# Glyphosate, a herbicide, and fosfomycin, an antibiotic in clinical use- evidence of common selectable genotypes

**DOI:** 10.64898/2026.05.15.725383

**Authors:** Katie D. Wall, Amy Campbell, Maitiú Marmion, Anna Kilroy, Caoimhe Doyle, Helina Marshall, Séamus Fanning

## Abstract

The emergence of antimicrobial resistance (AMR) is increasingly linked to metabolic adaptation, yet the evolutionary routes underlying cross-resistance between structurally related compounds remain poorly understood. Here, whole genome sequencing (WGS) was used to analyse *Klebsiella pneumoniae* mutants evolved under sub-lethal glyphosate (GLP) or fosfomycin (FOS) exposure to determine how these stresses shape resistance and physiology.

Sub-lethal GLP exposure increased FOS resistance, demonstrating cross-resistance between the two phosphonates. FOS-evolved mutants achieved high-level resistance through the accumulation of multiple mutations affecting the antibiotic target MurA, transport systems, and global metabolic regulation, producing a layered FOS resistance phenotype. In contrast, GLP-evolved mutants acquired similar functional classes of mutations but exhibited lower baseline FOS resistance, suggesting trade-offs between resistance and metabolic fitness. Further, analysis of FOS-evolved and GLP-evolved mutants across known bacterial GLP resistance mechanisms demonstrated a strong overlap.

Comparative genomic analysis revealed a small, recurrent set of genes under selection in both evolutionary trajectories, with identical genomic loci repeatedly targeted, consistent with convergent evolution. Many of these changes were linked to central metabolism, redox balance, and cell surface regulation. For some isolates, a hypermutator phenotype was necessary to offset the potentially lethal effects of primary-target mutations through compensatory genomic adaptation.

In conclusion, GLP and FOS select for shared adaptive networks that couple metabolic rewiring with AMR, revealing cross-resistance as an emergent property of global physiological reprogramming and providing mechanistic insight into ecological models of co-selection in environmental systems.

**Importance statement:** Glyphosate is a herbicide in current use worldwide. Its impact on the susceptibility of bacteria to antibiotics, remains to be described. This manuscript details a critical phenotypic and genomic analysis of shared resistance mechanisms between glyphosate and the antibiotic fosfomycin. Using *Klebsiella pneumoniae*, a zoonotic pathogen, this manuscript demonstrates that evolutionary adaptation to either compound results in a substantial overlap in gene mutations.

Crucially, this manuscript shows that exposure to sub-lethal concentrations of glyphosate can increase resistance to fosfomycin. These findings reveal a link between agricultural chemical use and the emergence of cross-resistance to fosfomycin. By highlighting how environmental factors drive antimicrobial resistance, this study underscores an urgent need for revision of food safety and regulatory frameworks to protect ***One Health***.

**Graphical abstract:** 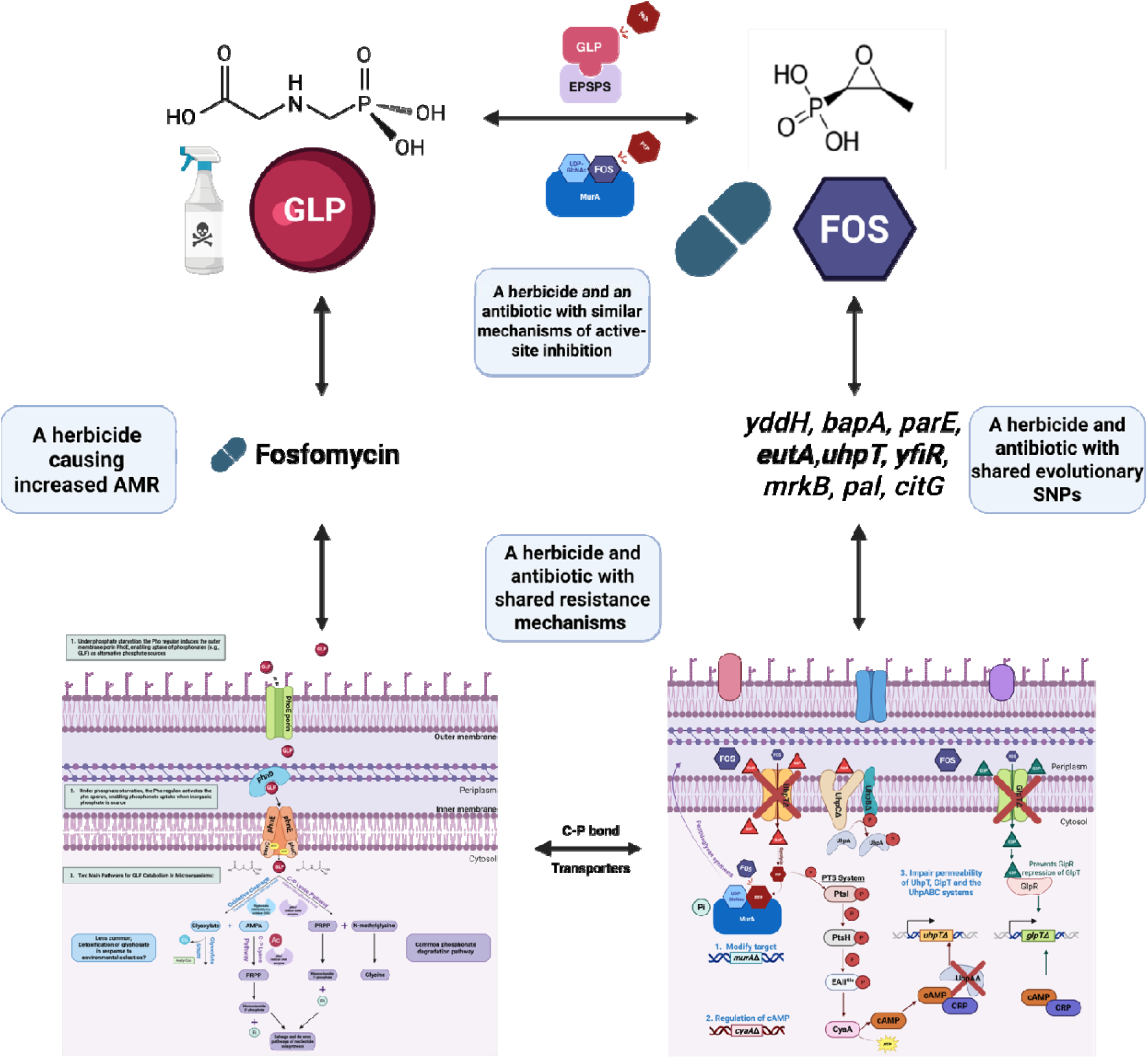

## Introduction

Increasing rates of antimicrobial resistance (AMR) in both Gram-positive and -negative pathogens, combined with a shortage of newly developed antimicrobial agents, have renewed interest in the use of older antibiotics. One such compound, Fosfomycin (FOS) [(1R,2S)-epoxypropylphosphonic acid], is an older bactericidal broad-spectrum antibiotic, historically used mainly for oral treatment of uncomplicated urinary tract infections. FOS has re-emerged as a promising option due to its retained activity against multidrug resistant (MDR) and extensively drug resistant (XDR) bacteria. Increasing evidence suggests that FOS remains effective against a broad range of resistant pathogens, prompting its reconsideration for *difficult-to-treat* infections (1). Although not widely used in animals, FOS represents a promising candidate for veterinary infections and has been used in the treatment of broiler chickens and pigs (2). The ***One Health*** relevance of retained efficacy of FOS against MDR and XDR pathogens in both human and animal settings underscores the need for coordinated stewardship across clinical and veterinary sectors to preserve its therapeutic value.

Peptidoglycan forms the rigid framework of the bacterial cell wall and, together with the outer membrane, plays a critical role in maintaining cellular integrity and preventing lysis under osmotic pressure (3, 4). Peptidoglycan synthesis **(Figure S1)** is mediated by two membrane transporters; <u>u</u>ptake <u>h</u>exose <u>p</u>hosphate <u>t</u>ransport, UhpT; which transports D-glucose-6-phosphate (G6P), and <u>gl</u>ycerol-3-<u>p</u>hosphate transporter, GlpT, which transports sn-glycerol-3-phosphate (G3P). UhpT is a major facilitator superfamily (MFS) transporter that exchanges inorganic phosphate (Pi) for sugar 6-phosphates (5). Under low Pi conditions, higher intracellular levels of G6P are required to initiate *uhpT* transcription due to inducer degradation mediated by the Pho regulon (6). Expression of *uhpT* is regulated by the UhpABC system (7).

In contrast, GlpT is a conserved organophosphate:phosphate antiporter of the MFS family (8). G3P is transported into the cell *via* GlpT in exchange for Pi, and acts as an inducer of the regulon by binding the transcriptional repressor GlpR. In the absence of G3P, GlpR represses *glpT* expression, whereas G3P binding relieves this repression, leading to increased *glpT* transcription (9, 10).

Under both aerobic and anaerobic conditions in *Enterobacteriaceae*, full expression of GlpT and UhpT requires elevated intracellular levels of adenosine 3′,5′-cyclic monophosphate (cAMP), which is synthesised by adenylate cyclase (CyaA; EC 4.6.1.1) and modulated by the phosphoenolpyruvate (PEP)-dependent phosphotransferase system (PTS) enzyme I (PtsI; EC 2.7.3.9). The global regulator cAMP- cAMP receptor protein (CRP) binds upstream of the *glpT* operon, counteracting repression by GlpR and promoting transcription. Similarly, cAMP-CRP binds to a site upstream of the *uhpT* promoter, where it acts in conjunction with the response regulator UhpA (11, 12).

Following its entry into central carbon metabolism, G6P can be metabolised *via* glycolysis to generate PEP, the high-energy phosphoryl donor used by the PTS to initiate phosphate transfer to PtsI. The phosphorylation state of EIIA^Glc^ reflects carbon availability and plays a central regulatory role. When EIIA^Glc^ is phosphorylated, it stimulates the activity of CyaA, leading to increased cAMP synthesis (13, 14). Alternatively, PEP serves as a substrate for MurA (UDP-*N*-acetylglucosamine enolpyruvyl transferase; EC 2.5.1.7), which transfers the enolpyruvate group from PEP to UDP-N-acetylglucosamine (UNAG), forming UDP-N-acetylglucosamine enolpyruvate (EP-UNAG) and releasing Pi in the initial step of peptidoglycan biosynthesis (15).

UhpT and GlpT permeases can also transport the antibiotic, FOS **(Figure S2)** (16). FOS is a PEP analogue that targets MurA. By binding to MurA, FOS blocks the formation of EP-UNAG from UNAG and PEP during the first committed step of cell wall synthesis, ultimately leading to bacterial lysis and cell death (3, 16). FOS inhibits MurA through covalent thioether bond formation with a conserved active-site cysteine (Cys_115_). Structural studies of the *E. coli* MurA-UNAG-FOS complex showed that the antibiotic is tightly positioned within the active site, where its phosphonate group forms strong electrostatic interactions with conserved positively charged residues (Lys22, Arg120, and Arg397) (17).

Bacteria can acquire resistance to FOS while maintaining peptidoglycan biosynthesis through three principal mechanisms **(Figure S3).** *First*, mutations or modifications of the target enzyme MurA can reduce or prevent FOS binding. *Second*, alterations in intracellular cAMP levels can diminish transcription of FOS uptake systems, which require cAMP for full expression. *Thirdly*, bacteria may limit antibiotic entry by impairing the function or permeability of the UhpT, GlpT, and UhpABC transport systems, thereby limiting intracellular FOS accumulation (18). In addition, enzymatic inactivation of FOS has been described in pathogenic bacteria and is mediated by one of three FOS-modifying enzymes, FosA [Mn(II) and K^+^-dependent glutathione transferase], FosB [Mg^2+^-dependent L-cysteine thiol transferase], or FosX [Mn(II)-dependent FOS-specific epoxide hydrolase], which chemically neutralise the antibiotic by cleaving its oxirane ring, rendering it inactive (19).

*N-(Phosphonomethyl)glycine*, commonly known as glyphosate (GLP), was first introduced commercially as a herbicide in 1974 and has since become the most widely used broad-spectrum herbicide worldwide (20). GLP, a PEP analogue, acts as a competitive inhibitor of 5-enolpyruvylshikimate-3-phosphate (EPSPS; EC 2.5.1.19), an enzyme encoded by the *aroA* gene in the shikimate pathway (21). The resulting aromatic amino acid (AAA) depletion causes plant death, and it similarly suppresses bacterial growth unless AAAs are externally supplied. This AAA deficiency disrupts multiple metabolic processes, explaining GLP’s pronounced impact on bacterial physiology (22, 23).

MurA (the target of FOS) and EPSPS (the target of GLP) are the only currently known PEP-dependent enolpyruvyl transferases (17). In addition to exhibiting similar modes of action, GLP and FOS are chemically related, as both belong to the phosphonic acid family and contain a stable Carbon-Phosphorous (C-P) bond **(Figure S4)** (24).

Notably, FOS resistance in both Gram-positive and -negative bacteria isolated from humans and animals appears to have increased over the past four decades. Several studies reported particularly high rates of FOS resistance in regions with extensive GLP usage, including China, raising the possibility of an association between widespread GLP exposure and elevated FOS resistance (25, 26). Given the shared chemical properties of FOS and GLP, as well as their convergence on PEP-dependent pathways, it is plausible that resistance mechanisms to these compounds may intersect at the level of bacterial metabolism. This potential overlap represents a critical interface within the ***One Health*** paradigm, linking environmental GLP exposure with AMR in animal and human pathogens. While a direct causal relationship has not been established, the possibility that widespread GLP use contributes indirectly to FOS resistance warrants further investigation.

In this study, WGS of FOS- and GLP-evolved *Klebsiella* mutants, together with phenotypic antimicrobial susceptibility testing for FOS in the presence and absence of sub-lethal GLP, showed that combined exposure to GLP and FOS selects for interconnected adaptive pathways that promote AMR. Analysis of shared SNPs, and those mapped to known resistance mechanisms for GLP and FOS, revealed overlapping genetic determinants associated with resistance, virulence, and metabolism. Together, these findings show that environmental chemical exposure can drive unintended and complex adaptive responses that extend beyond resistance to the selecting agent itself, with important implications for the emergence and persistence of MDR.

## Materials and methods

### Bacterial isolates

*Klebsiella pneumoniae*: MGH 78578 (ATCC^TM^700721), was cultured on Columbia agar (Oxoid, CM0331) with 5% [v/v] defibrinated horse blood (TCS Biosciences, HB034) (CBA) for 24 hours at 37°C being taken from stocks stored at -80°C at the UCD-Centre for Food Safety.

### Determination of minimum inhibitory concentration (MIC) by antibiotic broth microdilution

The MIC of the type strain *K. pneumoniae* MGH 78578 was tested against glyphosate (Sigma Aldrich, 89432), and fosfomycin (Sigma Aldrich, P5396) supplemented with 25 mg/L glucose-6-phosphate (Sigma Aldrich, 346764), as mandated by EUCAST when obtaining a fosfomycin MIC (EUCAST. [(accessed on 11^th^ May, 2026)]; Clinical Breakpoints. Available online: http://www.eucast.org/clinical_breakpoints). The MIC was determined according to the method previously described (27) with the use of Brucella broth (Becton, Dickinson and Company, 211088. This change was made as cations interfere with GLP activity (28). Sub-inhibitory concentrations of GLP were further used in the study at 0.25 X the MIC of GLP **(Figure S5)**. All the tests were performed in three biological replicates unless stated otherwise.

### Generation of isogenic glyphosate and fosfomycin evolutionary mutants

The same method was used for generating the GLP and FOS-evolved mutants. For the preadaptation passage, single colonies of *K. pneumoniae* MGH 78578 were isolated from CBA and passaged daily 1:100 in 2 mL Brucella broth (as per the guidelines above, G6P was added to the broth for the FOS evolution) for 3 days to help bacteria to adapt to the experimental conditions **(Figure S6)**. For exposure, the starting subinhibitory concentration for the challenge was 0.25X of the obtained MIC for the active (GLP or FOS). Each day, 20 μL of the overnight culture was transferred into two new tubes, one with the same concentration of the active at which the visible growth was observed in the last passage and one with a concentration of the active 0.25X greater again than the last. Nonevolving controls were handled similarly with the exception that the active was absent. Serial passages were terminated once no visible growth was detected at the subsequent higher concentration. For WGS, mutant lineages were harvested and cryopreserved at various inhibitory concentrations to capture diverse resistance profiles. This protocol was adapted from (29).

### DNA template preparation and whole genome sequencing

All study isolates **(Table 1)** were cultured in 10 mL Tryptone Soya Broth (Oxoid, CM0129) overnight (for 18 h) at 37°C and were used as starting material for the DNA purification using the DNeasy UltraClean Microbial Kit (Qiagen, 10196-4) according to manufacturer’s instructions and the protocol for Gram-negative bacteria. DNA concentration and quality was assessed using the Qubit fluorometer (Thermo Fisher Scientific) and Tapestation platform (Agilent). Each DNA concentration was normalised to 500 ng in 40 μL and used as starting template for library preparation with the Illumina DNA preparation kit [Illumina, DNA Prep: IPB and Buffers (20049006), PCR and Buffers (20015829, DNA/RNA UD Indexes Set A (200916460) and Set B (20091647), and MiSeq reagent kit V3 (600 cycles) (MS-102-3003) following the manufacturer’s instructions to obtain a final library concentration of 4 nM. The final denatured DNA libraries at 12 pM concentrations were combined with 5% PhiX (Illumina, 15017397) and loaded using the MiSeq Reagent Kit v3 (600-cycle) (Illumina, MS-102-3003). Data was collected for 76 hours. DNA sequencing was performed inhouse.

**Table 1.** Kleborate and Pathogenwatch outputs for the GLP and FOS evolutionary *Klebsiella* mutants, compared to the isogenic control *Kp*_type.

| *Klebsiella* | ST | cgMLST | SL | CG | Inc types | K locus | O locus | Resistance score | Virulence score |
| --- | --- | --- | --- | --- | --- | --- | --- | --- | --- |
| *Kp_type* | 38 | 25330 | 38 | 38 | Col440I;<br>IncFIB(K);<br>IncFIB(pQII);<br>IncFII(K);<br>IncR | KL52 | OL13 | 0 | 0 |
| *Kp_GLPMUT1* | 38 | 25330 | 38 | 38 | IncFIB(pQII);<br>IncFII(K);<br>IncR | KL52 | OL13 | 1 | 0 |
| *Kp_GLPMUT2* | 38-2LV | *a165 | New | New | IncFIB(pQII);<br>IncFII(K);<br>IncR | KL52 | OL13 | 0 | 0 |
| *Kp_GLPMUT3* | 38 | *1455 | 38 | 38 | IncFIB(pQII);<br>IncFII(K);<br>IncR | KL52 | OL13 | 0 | 0 |
| *Kp_FOSMUT1* | 38 | 25330 | 38 | 38 | Col440I;<br>IncFIB(K);<br>IncFIB(pQII);<br>IncFII(K);<br>IncR | KL52 | OL13 | 0 | 0 |
| *Kp_FOSMUT2* | 38 | 25330 | 38 | 38 | Col440I;<br>IncFIB(K);<br>IncFIB(pQII);<br>IncFII(K);<br>IncR | KL52 | OL13 | 1 | 0 |
| *Kp_FOSMUT3* | 38 | *7494 | New | New | Col440I;<br>IncFIB(K);<br>IncFIB(pQII);<br>IncFII(K);<br>IncR | KL52 | OL13 | 0 | 0 |
| *Kp_FOSMUT4* | 38 | 25330 | 38 | 38 | Col440I;<br>IncFIB(K);<br>IncFIB(pQII);<br>IncFII(K);<br>IncR | KL52 | OL13 | 0 | 0 |
| *Kp_FOSMUT5* | 38 | 25330 | 38 | 38 | Col440I;<br>IncFIB(K);<br>IncFIB(pQII);<br>IncFII(K);<br>IncR | KL52 | OL13 | 0 | 0 |
| *Kp_FOSMUT6* | 38 | 25330 | 38 | 38 | IncFIB(K);<br>IncFIB(pQII);<br>IncFII(K);<br>IncR | KL52 | OL13 | 0 | 0 |
| *Kp_FOSMUT7* | 38 | 25330 | 38 | 38 | Col440I;<br>IncFIB(K);<br>IncFIB(pQII);<br>IncFII(K);<br>IncR | KL52 | OL13 | 1 | 0 |

Quality control of raw sequencing reads was first assessed using FastQC (Galaxy *version 1.20.0;* (112)). Reads were then filtered and trimmed, where necessary, using FastP (Galaxy *version 1.0*.*1;* (30)) to ensure a minimum Phred quality score of 30 across all reads. High quality reads were assembled using Shovill, a SPAdes-based genome assembler (Galaxy *version 1.1.0;* (113)). Assembly graphs were inspected using Bandage Info and Bandage Image (Galaxy *version 2022.09;* (31)), and assembly quality was evaluated with QUAST (Galaxy *version 5.3.0*); (32).

Taxonomic profiling for species identification and assessment of potential contamination was performed using Kraken2 (Galaxy *version 2.1.3;* (33)), Bracken (Galaxy *version 2.9;* (34)), and Recentrifuge (Galaxy *version 1.14.0;* (*35*)). Genomes were further analysed for sequence type, K-type, O-type and the presence of resistance and virulence determinants using Kleborate (Galaxy *version 2.3.2*) (36) and Pathogenwatch (https://pathogen.watch *version 23.5.0*). Antimicrobial resistance and plasmid content were assessed using ABRicate (Galaxy *version 1.0.1;* (114)) with the CARD (Database version: 2020-April-19; (37)) and PlasmidFinder (Database version: 2020-April-19; (38)) databases and results were summarised using ABRicate Summary (Galaxy v*ersion 1.0.1;* (114)).

Genome annotation was carried out using Prokka (Galaxy *version 1.14.6;* (39)), and pangenome analysis was performed with Roary ((Galaxy *version 3.13.0*) (40). Phylogenetic and downstream analyses were conducted in RStudio (*version 2024.04.2+764;* (115)), with tree visualisation performed using iTOL interactive tree of life (*version 7.3)* (41).

Single nucleotide polymorphism (SNP) analysis was conducted using Snippy and snippy-core (Galaxy *version 4.6.0;* (116)), and pairwise SNP distances were calculated using the SNP distance matrix tool ((Galaxy *version 0.8.2;* (117)). Graphs were created in BioRender (https://BioRender.com).

### Fosfomycin e-Strip agar method

According to EUCAST, agar dilution is the reference method for obtaining the MIC to FOS **(Figure S5)**. All study isolates were grown on CBA, and Fosfomycin MIC Test Strips 0.064-1,024 μg/mL (the strips contained G6P; Liofilchem, 920790) were used to test susceptibility according to Liofilchem^®^- Fosfomycin MIC Test Strip Technical Sheet - MTS45 - Rev.1 / 09.11.2017.

A 0.5 McFarland standard was obtained and isolates streaked on unmodified CBA and a FOS MIC test strip applied (control plate [CP]), and CBA dissolved with 0.25X GLP and a FOS MIC test strip applied (test plate [TP]). The plates were incubated overnight at 37°C and both the CP and TP were read visually after 20 hours and compared to the EUCAST breakpoint for *Eubacterales* (Version 13.0) for the determination of the MIC. All the tests were performed in three biological replicates unless stated otherwise.

## Results & Discussion

### MIC measurements and WGS-based assessment of GLP- and FOS- induced genome changes in *Klebsiella*

In *K. pneumoniae* MGH 78578 (denoted as *Kp*_type/ST38) the MIC to GLP was determined to be 3 mg/mL. Exposure of *Kp_type* to increasing concentrations of GLP resulted in the selection of three isogenic GLP-evolved mutants (**Table 1**; and **Table S1**). These were denoted as *Kp_GLPMUT1* and *Kp_GLPMUT2* and exhibited growth up to 1.9X the MIC GLP (5.7 mg/mL), while in contrast *Kp_GLPMUT3* grew up to 2X the MIC (6 mg/mL) **(Figure S5)**. Both *Kp_GLPMUT1* and *Kp_GLPMUT3* retained the same sequence type (ST38) as the isogenic control and acquired 38 and 51 single nucleotide polymorphisms (SNP), respectively. In contrast, *Kp_GLPMUT2* exhibited a markedly higher number of SNPs (n = 6,056) and was assigned to ST38-2LV, indicating it differed at two loci (conserved housekeeping genes) from ST38 based on the MLST scheme for *K. pneumoniae* and related species.

In *Kp*_type, the MIC against FOS **(Table S3)** was determined by broth microdilution to be 80 µg/mL across three biological replicates. This MIC was used as the baseline for the generation of FOS-evolved *Klebsiella* mutants (**Table 1)**. Stepwise exposure of the parental *Kp*_type strain to increasing concentrations of FOS resulted in the selection of seven isogenic FOS-evolved mutants, with the most resistant isolate, *Kp*_FOSMUT7, able to grow at concentrations up to five times the MIC (400 µg/mL) **(Table S2;** and **Table S3).** All seven FOS-evolved mutants retained sequence type ST38.

WGS analysis carried out after selection on GLP, showed that all three GLP-evolved mutants lost the Col440I and IncFIB(K) replicon type plasmids, a feature that could be expected to reduce fitness burden **(Table 1)**. Col440I plasmids are typically small, while IncFIB(K) plasmids are larger and both are commonly found in *Enterobacteriaceae* (42). PlasmidFinder also detected a ColRNAI plasmid in *Kp_type* and the three GLP evolutionary mutants; when co-carried with IncR, this plasmid combination has been linked to multidrug-hypervirulence (MDR-hv) in ST395 in *K. pneumoniae* (43). In contrast, plasmid loss was observed in only one FOS-evolved mutant, *Kp*_FOSMUT6, wherein it lost the small Col440I plasmid.

*Kp*_GLPMUT1, *Kp*_FOSMUT2, and *Kp*_FOSMUT7 carried a non-wildtype SHV beta-lactamase, SHV-12, a well characterised ESBL variant which resulted in a Kleborate resistance score of 1. The SHV-12 contained three key amino acid substitutions (denoted as 238S;240K;35Q), associated with the evolution of ESBL activity from parental SHV enzymes such as SHV-5 or SHV-11 (44). These substitutions expand substrate specificity, conferring resistance to third-generation cephalosporins (45). In *Klebsiella*, ESBL-encoding *bla*_SHV_ genes are most often plasmid-borne, and SHV-12 is frequently associated with IS*26*-mediated transposons, which facilitate mobilisation and dissemination (46).

### Comparative analysis of SNPs in *Kp*_GLPMUT2 and *Kp*_FOSMUT3

Analysis of the pangenome for the *Kp*_type strain, the three GLP-evolved mutants and the seven FOS-evolved mutants (**Table S4**) revealed a total of 5,627 genes shared among the *Kp*_type and its evolved mutants, comprising 4,897 core genes, 366 shell genes, and 364 cloud genes.

*Kp*_GLPMUT1, *Kp*_GLPMUT3, *Kp*_FOSMUT1, *Kp*_FOSMUT2, *Kp*_FOSMUT4, *Kp*_FOSMUT5, *Kp*_FOSMUT6, and *Kp*_FOSMUT7 were phylogenetically similar to the parental *Kp*_type, differing by 38, 51, 11, 14, 22, 13, 14, and 21 SNPs respectively. In contrast, *Kp*_GLPMUT2 and *Kp*_FOSMUT3 were markedly divergent, harbouring 6,056 and 7,317 SNPs, respectively, relative to *Kp*_type, and differing by 13,188 SNPs from each other (**Figure 1**). Genomic analysis revealed that the high SNP density was driven by a hypermutator phenotype resulting from the loss of primary DNA maintenance pathways (**Table S5**). In this context, neutral passenger mutations likely co-segregated with beneficial alleles, providing the requisite genetic variation to facilitate adaptation to both GLP and FOS. Most notably, *Kp*_GLPMUT2 harboured a premature stop codon in ***dnaQ*** (Lys32*). As *dnaQ* encodes the epsilon subunit responsible for the 3’-to-5’ exonuclease activity of DNA Polymerase III, this truncation effectively abolishes the replicative proofreading capability of the cell (47). Both highly mutated lineages showed significant hits to the MutHLS system in the mismatch repair (MMR) pathway. *Kp*_GLPMUT2 carried a stop-gain in ***mutS*** (Gln476*), missense variants in ***mutL*** (Aly515Glu;Lys307Thr), and a synonymous variant in **MutH** (Leu78Leu), while *Kp*_FOSMUT3 harboured missense mutations in ***mutS*** (leading to a Ile279Val amino acid substitution) and similarly in ***mutL*** (His308Tyr). Further compounding the repair deficit were mutations identified in DNA Helicase II ***uvrD,*** which is essential for unwinding the DNA strand during excision repair (48). *Kp*_GLPMUT2 had a stop-gain (Glu447*) and missense variants in *uvrD* (Asp250Glu;Asp311Ala), as did Kp_FOSMUT3 (Arg375His;Val720Gly).

**Figure 1.**
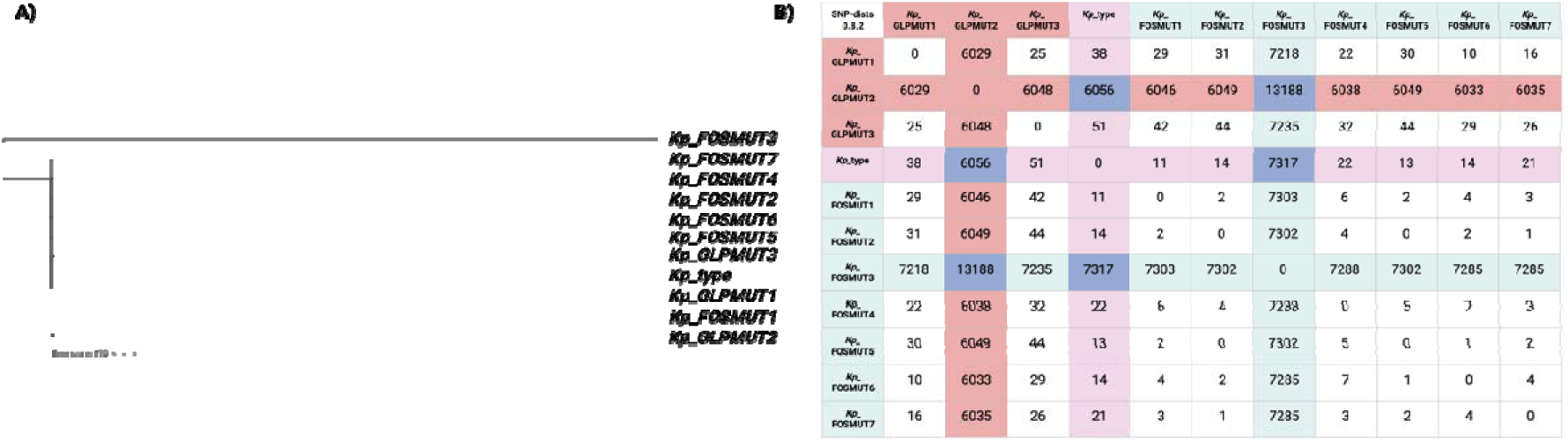
Genomic divergence and phylogenetic relationships between *Kp*_type and its GLP- and FOS-evolved mutants. **A)** a phylogenetic tree including the isogenic control *Kp*_type strain and its three GLP-evolved and seven FOS-evolved mutants. The phylogenetic tree was created using iTOL. **B)** a summary of pairwise SNP distances, calculated using the SNP distance matrix tool.

In addition to the primary mismatch repair deficits, *Kp*_GLPMUT2 and *Kp*_FOSMUT3 harboured mutations in the ‘GO system’ (*mutT* and *mutY*), which is responsible for mitigating oxidative DNA damage. Given that both GLP and antibiotic stress are associated with the production of reactive oxygen species (ROS) (49, 50), the impairment of the 8-oxo-G repair pathway likely along with the MMR deficiency to drive the high frequency of transversion mutations observed in these lineages.

The link between defective DNA repair and chemical pressure suggests that *K. pneumoniae* utilises hypermutation as a critical evolutionary strategy to survive potent selective pressures. While other mutants (*Kp*_GLPMUT1/3 and *Kp*_FOSMUT1,2,4,5,6,7) bypassed traditional pathways via alternative resistance mechanisms **(Figures -1 & -2)**, *Kp*_GLPMUT2 and *Kp*_FOSMUT3 that acquired SNPs in primary targets (*aroA* and *murA* **(Figures -4, -5 & -6)**) developed a hypermutator phenotype. This suggests that because primary-target mutations can be potentially lethal or carry extreme fitness costs, a hypermutator state is required to facilitate the rapid acquisition of compensatory genetic changes necessary for survival.

Accordingly, a comparative analysis was performed focusing on gene-level SNPs identified in *Kp*_GLPMUT1, *Kp*_GLPMUT3, *Kp*_FOSMUT1, *Kp*_FOSMUT2, *Kp*_FOSMUT4, *Kp*_FOSMUT5, *Kp*_FOSMUT6, and *Kp*_FOSMUT7, followed by assessment of whether these SNPs were also present in the hypermutated *Kp*_GLPMUT2 and *Kp*_FOSMUT3 **(Figure 2).** This analysis also aimed to identify any shared SNPs common to all evolutionary mutants.

**Figure 2.**
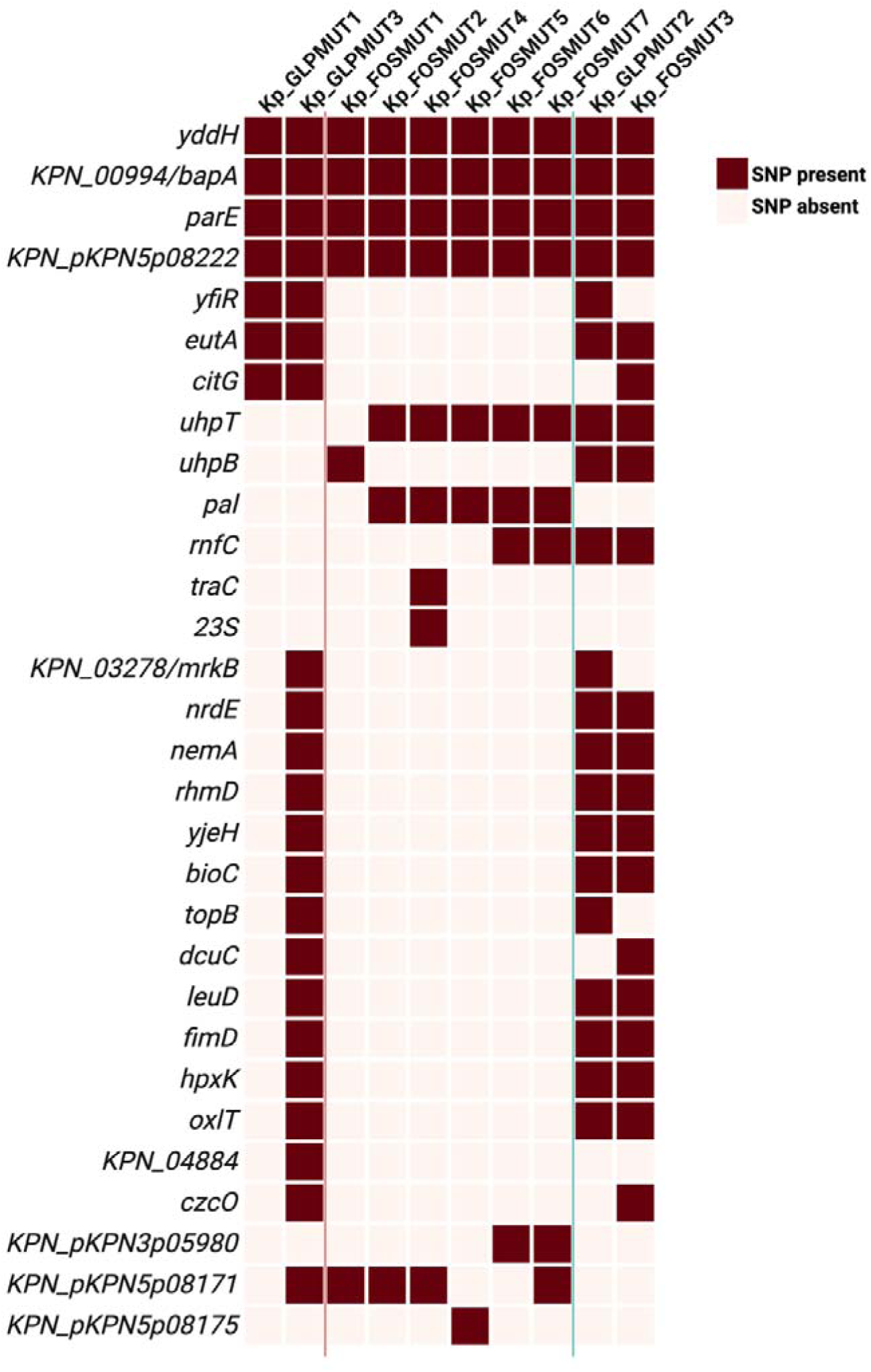
A comparison of SNPs identified in *Kp*_GLPMUT1, *Kp*_GLPMUT3, *Kp*_FOSMUT1, *Kp*_FOSMUT2, *Kp*_FOSMUT4, *Kp*_FOSMUT5, *Kp*_FOSMUT6, and *Kp*_FOSMUT7, and assessment of their presence in the hypermutated *Kp*_GLPMUT2 and *Kp*_FOSMUT3. The pink vertical line separates *Kp*_GLPMUT1 and *Kp*_GLPMUT3 from *Kp*_FOSMUT1, *Kp*_FOSMUT2, *Kp*_FOSMUT4, *Kp*_FOSMUT5, *Kp*_FOSMUT6, and *Kp*_FOSMUT7. The green vertical line further separates *Kp*_GLPMUT2 and *Kp*_FOSMUT3. *To note, the full names of the other genes containing SNPs are described in **Table S6**.

Comparative SNP analysis across all ten evolutionary mutants revealed four genes that were consistently altered in all strains. These included *yddH* (KPN_01888), encoding a flavin reductase family protein; KPN_00994, encoding a BapA prefix-like domain-containing protein; *parE*, encoding DNA topoisomerase IV subunit B; and KPN_pKPN5p08222, encoding a hypothetical protein located on plasmid pKPN5.

Strikingly, all ten mutants harboured the same missense variants in ***yddH***, namely **Arg114Leu** and **Ala118Glu**. *Kp*_FOSMUT3 contained an additional *yddH* variant (Asp152Glu). However there is limited literature describing the functional role of *yddH*, and its contribution to bacterial physiology remains poorly characterised.

One well-characterised role of reduced flavins is in the activation of ribonucleotide reductase, where they facilitate reduction of the Fe^3+^ centre and subsequent formation of a catalytically essential tyrosyl radical (51). Given this important role in redox balance and DNA synthesis, mutations affecting flavin reductase-like genes such as *yddH* may indirectly influence cellular energy status and stress responses. As the ferrous iron (Fe^2+^) participates in the Fenton reaction to produce highly reactive hydroxyl radicals, bacteria tightly regulate iron redox homeostasis as part of their oxidative stress response. Flavin reductases have been implicated in iron redox balance and oxidative stress defence by influencing Fe^2+^ pools and limiting ROS-mediated damage (52, 53).

The consistent acquisition of identical *yddH* missense mutations across all ten evolutionary mutants strongly suggests positive selection acting on this locus. Given the predicted flavin reductase-like function of YddH, these substitutions are likely to modulate cellular redox balance. By subtly altering flavin-mediated electron transfer, the *yddH* variants may reduce intracellular Fe^2+^availability, thereby limiting Fenton-mediated ROS formation under chemical stress. Such a shift in iron redox homeostasis could enhance tolerance to oxidative and antibiotic-associated stress by promoting a more reductive intracellular environment.

KPN_00994 encodes a **BapA prefix-like domain-containing protein** comprising 2,744 AA residues. Owing to its large size, numerous SNPs were identified within this gene across the ten evolutionary mutants, occurring at diverse positions. For example, *Kp*_GLPMUT2 alone contained 14 SNPs within KPN_00994. In *Acinetobacter baumannii*, the biofilm-associated protein (BAP) is a large surface protein that contributes to biofilm formation and adhesion (54). Carbapenem-resistant *A. baumannii* (CRAB) is a major cause of hospital outbreaks worldwide. Genomic diversification during the outbreak was driven primarily by recombination and localised sequence variation at specific loci in two circulating clones. Notably, both clones exhibited a high density of SNPs in the BapA protein, implicating this adhesin as a key target of adaptive evolution linked to biofilm persistence and host-environment interactions during outbreak spread (55). Taken together, the extensive and distributed SNP accumulation observed in KPN_00994 across the evolutionary mutants is consistent with patterns reported for large BapA-like biofilm-associated proteins, which frequently act as mutational hotspots during adaptive evolution.

All ten mutants harboured the same missense variant in ***parE***, **Tyr4Asp**. *Kp*_FOSMUT3 contained an additional *parE* variant (Thr32Ser), as did *Kp*_GLPMUT2 (Gly131Gly; Asp197Val; Leu280Leu). DNA gyrase and topoisomerase IV are heterotetrameric enzymes composed of GyrA/GyrB and ParC/ParE subunits, respectively. While the GyrA and ParC subunits mediate DNA binding and catalysis, GyrB and ParE provide the ATP-binding activity; consequently, ParE (topoisomerase IV subunit B) is essential for the energy-transducing function and covalent catalysis required for topoisomerase IV activity (56, 57). Owing to its close homology with *gyrB*, *parE* contains a conserved quinolone resistance-determining region QRDR. In ParE, this region (Asp420-Lys441) corresponds directly to the QRDR of GyrB (Asp426-Lys447) (58). Collectively, these findings suggest that the recurrent *parE* Tyr4Asp substitution is unlikely to represent a canonical quinolone-resistance mutation, as it lies outside the established ParE QRDR and *parE* mutations typically contribute to resistance only after primary *gyrA* (with and without *parC*) alterations have reduced gyrase susceptibility. Instead, the convergence on the same ParE substitution across all mutants may reflect selection for an indirect fitness advantage, potentially through altered topoisomerase IV ATPase activity or replication-associated stress tolerance. Additional ParE substitutions observed in the hypermutated lineages may further modulate enzyme function.

All three evolutionary GLP mutants, as well as *Kp*_FOSMUT3, harboured SNPs in ***eutA***, which encodes the ethanolamine ammonia-lyase (EAL) reactivase. Similarly, all three GLP-evolved mutants also shared a synonymous variant (**Ile433Ile)**. In addition, *Kp*_GLPMUT2 carried a nonsynonymous variant (His191Gln), while *Kp*_FOSMUT3 contained five SNPs within *eutA* (Arg224Leu; Gly187Gly; Thr139Ala; Val129Val; Asp58Tyr).

EAL catalyses the deamination of ethanolamine to acetaldehyde and ammonia, with subsequent dismutation of acetaldehyde yielding acetate and ethanol (59). Phosphatidylethanolamine (PE) is a major phospholipid component of both bacterial and mammalian cell membranes and can be degraded by phosphodiesterases to yield ethanolamine and glycerol-derived products (60, 61). Ethanolamine can then be utilised as a carbon and/or nitrogen source following cleavage by EAL within a bacterial microcompartment, which functions to sequester the volatile intermediate acetaldehyde (62). Acetaldehyde may subsequently be reduced to ethanol by EutG (alcohol dehydrogenase; EC 1.1.1.1) or converted to acetyl-CoA by EutE (acetaldehyde dehydrogenase; EC 1.2.1.10) (63), allowing entry into central metabolic pathways including the tricarboxylic acid cycle (TCA) cycle, the glyoxylate shunt, and lipid biosynthesis (64). Alternatively, acetyl-CoA may be channelled through EutD (phosphotransacetylase; EC 2.3.1.8) to form acetyl phosphate, which is then converted to acetate by the housekeeping acetate kinase AckA (EC 2.7.2.1), yielding ATP (65). These changes may also reflect an adaptive strategy to preserve membrane integrity under stress by limiting PE turnover. As ethanolamine metabolism is frequently exploited by enteric pathogens under anaerobic gut conditions to confer a competitive advantage (66, 67), its potential suppression in these mutants suggests a metabolic shift away from host-associated nutrient utilisation toward alternative survival or stress-adaptive strategies. This may reflect reduced reliance on ethanolamine-derived acetyl-CoA feeding into the TCA cycle and glyoxylate shunt, potentially favouring glycerol utilisation or other fermentative metabolic pathways that impose lower demands on NADH generation. In *Salmonella enterica*, coordinated regulation of the *eut* and *pdu* operons prevents interference between their structurally similar microcompartments, with 1,2-propanediol inducing the *pdu* operon while simultaneously repressing the *eut* operon to avoid deleterious mixing of the two systems (68).

All evolutionary FOS mutants and *Kp*_GLPMUT2 harboured SNPs in ***uhpT***, with the exception of *Kp*_FOSMUT1, which instead carried a SNP in ***uhpB***. *Kp*_GLPMUT2 and *Kp*_FOSMUT3 also carried *uhpB* SNPs. The *uhpB* SNP in *Kp*_FOSMUT1 is a disruptive in-frame deletion (*c.*1293_1298delCAGCGC) resulting in the loss of Ser432 and Ala433 (*p.*Ser432_Ala433del). All evolutionary FOS mutants harboured the same *uhpT* missense variant, **Arg46His**. The biological implications of *uhpT* mutations are discussed in detail below, but together these findings further support the conclusion that blocking the G6P importer is critical for GLP and FOS resistance.

All three evolutionary GLP mutants harboured SNPs in *yfiR*, which encodes a YfiR family protein. *Kp*_GLPMUT1 and *Kp*_GLPMUT2 shared a missense substitution (**Arg6Leu**), while *Kp*_GLPMUT2 carried an additional mutation (**Cys89Ser**) and *Kp*_GLPMUT3 contained a distinct substitution (**Thr40Pro**). The YfiBNR signalling system is a periplasmic regulatory module that controls intracellular bis(3′-5′)-cyclic diguanosine monophosphate (cyclic di-GMP) levels in Gram-negative bacteria. The Arg6Leu substitution lies within the predicted signal peptide of YfiR. Disruption of YfiR localisation has been shown to impair its ability to repress the diguanylate cyclase YfiN, leading to elevated intracellular cyclic di-GMP levels. The Thr40Pro substitution occurs within the N-terminal periplasmic region of the mature YfiR and may affect local protein folding or stability, while Cys89Ser alters a cysteine residue that lies within the periplasmic domain. Although Cys89 is not among the four cysteines strictly conserved in *P. aeruginosa* YfiR (two disulfide bonds Cys71-Cys110 and Cys145-Cys152), substitution of periplasmic cysteines has been associated with YfiR misfolding and loss of repression, potentially through altered disulfide bond formation or redox sensitivity (69, 70). In this system, YfiR functions as a periplasmic repressor of YfiN, while YfiB (an OmpA/Pal-like outer-membrane lipoprotein) relieves this repression under envelope or stress signals. Loss or impairment of YfiR activity results in derepression of YfiN, increased cyclic-di-GMP, reduced motility, and enhanced biofilm formation, phenotypes strongly associated with persistence and antibiotic tolerance (71).

In *K. pneumoniae*, type 3 fimbriae expression and biofilm formation are controlled by a cyclic-di-GMP-dependent regulatory network in which the transcriptional activator MrkH responds to intracellular c-di-GMP to stimulate *mrkABCDF* expression. Both MrkJ and YfiN, in *K. pneumoniae*, coordinate intracellular c-di-GMP concentrations to control MrkH-dependent regulation (72). Consistent with this, SNPs were also identified in ***mrkB*** in *Kp*_GLPMUT2 and *Kp*_GLPMUT3 (Ser160Thr and Phe10Leu respectively), suggesting coordinated selection of adhesion and surface-associated traits. Notably, FOS-evolved mutants (with the exception of *Kp*_FOSMUT1 and *Kp*_FOSMUT3) did not harbour *yfiR* mutations, but instead contained the same SNP encoding the peptidoglycan-associated lipoprotein (***pal)*** (**Thr27Pro**). Pal shares structural homology with YfiB and they both contain a peptidoglycan-binding domain (73), indicating that distinct evolutionary routes may converge on envelope-associated signalling and surface remodelling, either through YfiBNR-mediated regulation or through alterations in cell-envelope scaffolding proteins.

*Kp*_GLPMUT1, *Kp*_GLPMUT3, and *Kp*_FOSMUT3 harboured the same missense variant in ***citG***, encoding a 2-(5’’-triphosphoribosyl)-3’-dephosphocoenzyme-A synthase (EC 2.4.2.52), **Trp149Arg**. *Kp*_FOSMUT3 contained an additional *citG* variant (Arg147Arg). CitG catalyses an unusual phosphoribosyl transfer reaction in which ATP and dephospho-CoA are converted to 2′-(5″-triphosphoribosyl)-3′-dephospho-CoA, releasing free adenine as a by-product. This reaction initiates biosynthesis of the citrate lyase prosthetic group and is essential for citrate fermentation (74). *K. pneumoniae* ferments citrate anaerobically *via* dedicated *cit* operons whose expression is tightly controlled by catabolite repression through the cAMP-CRP regulatory system, ensuring that citrate utilisation is activated only when preferred carbon sources such as glucose, gluconate, and glycerol, are unavailable (75).

Together with suppression of ethanolamine utilisation, these findings suggest a metabolic shift away from host-derived ethanolamine toward alternative carbon sources, with glycerol remaining a likely preferred substrate before induction of citrate fermentation under catabolite repression control. The repeated inactivation of *uhpT* (G6P importer) but not *glpT* (G3P importer) further supports a shift away from host-derived phosphorylated substrates toward glycerol-associated metabolism, while simultaneously conferring resistance to GLP and FOS.

When considered together, these data indicate that adaptation to GLP and FOS selection pressures in *K. pneumoniae* involves integrated modulation of redox-related processes, cell-envelope and biofilm-associated signalling networks, and central metabolism, collectively favouring persistence, stress tolerance, and metabolic flexibility over maximal growth efficiency.

### Assessment of GLP-induced FOS resistance in *Klebsiella*

For the *Kp_type* strain, the addition of a sub-lethal concentration of GLP increased the MIC_FOS_ to 230 µg/mL, as determined by broth microdilution. This represents a nearly three-fold increase over the baseline MIC_FOS_ of 80 µg/mL used to initiate the FOS-evolution experiments (**Table S3**).

MIC values for all isolates were further determined for FOS in the presence and absence of a sub-inhibitory concentration of GLP by agar dilution and e-Strips, as recommended by EUCAST **(Figure 3)**. The FOS-evolved mutants were determined to have an MIC of 5 mg/mL to GLP **(Figure S5)**. ABRicate identified the FOS resistance gene *fosA6* in all assemblies, employing the CARD database. At present, there is limited clinical evidence to support established EUCAST breakpoints for FOS in *Enterobacterales*. However, for reference, the EUCAST susceptibility breakpoint for FOS in *E. coli* associated with urinary tract infections is 8 mg/L, and this was used in this study for reference breakpoint purposes.

**Figure 3.**
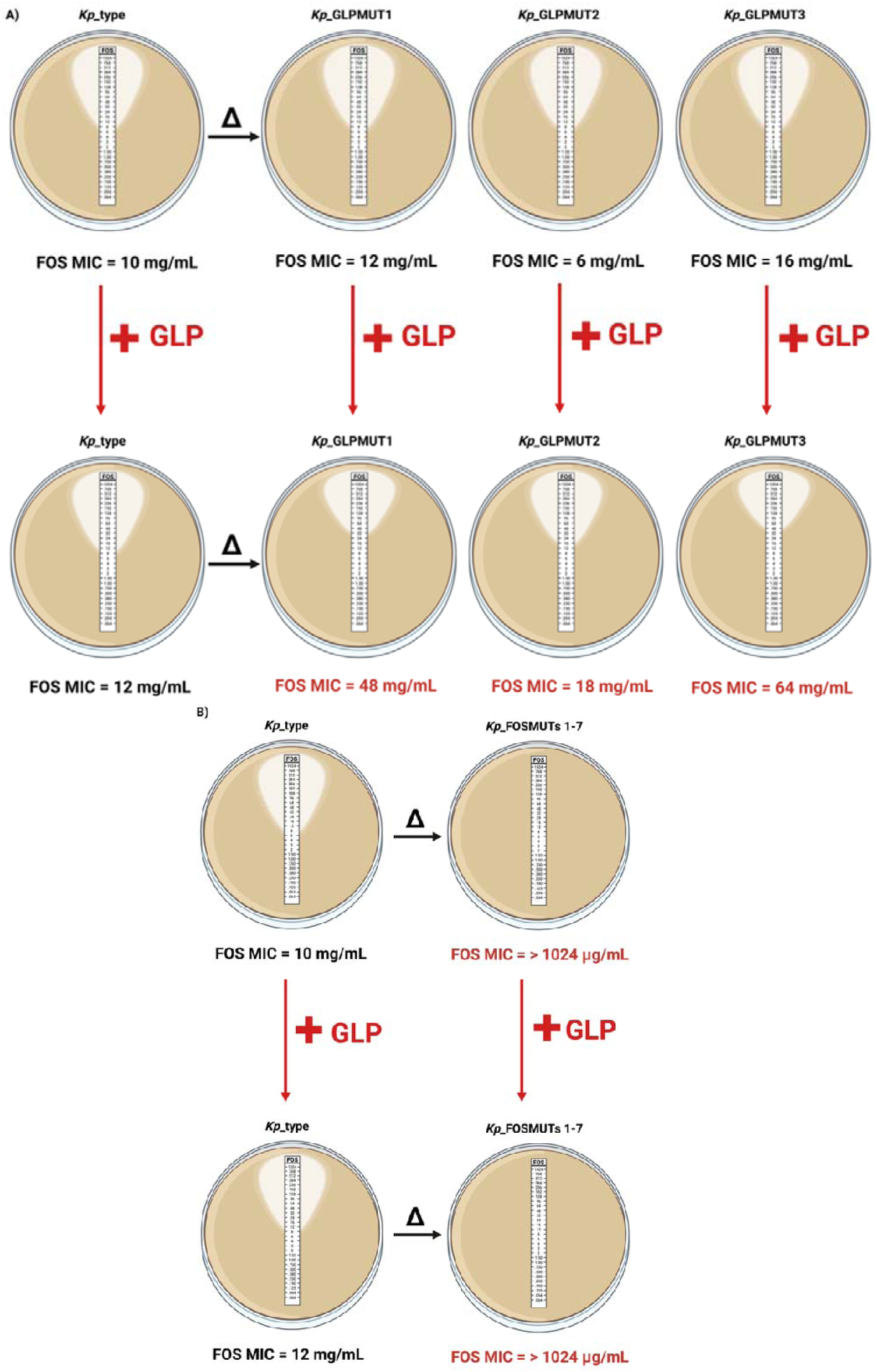
Sub-lethal GLP exposure alters antimicrobial susceptibility profiles to FOS in *Klebsiella*. MICs (μg/mL) to FOS are shown for the isogenic control *Kp*_type strain and **A)** its three GLP-evolved (GLPMUT1-3) and **B)** seven FOS-evolved (FOSMUT1-7) mutants, with and without 0.25X MIC GLP exposure.

In the case of the *Kp*_type isolate, exposure to GLP increased the FOS MIC from 10- to 12-µg/mL. The baseline MICs for *Kp*_GLPMUT1 and *Kp*_GLPMUT3 were already higher than that of *Kp*_type, at 12 µg/mL and 16 µg/mL, respectively. In the presence of GLP, their FOS MICs increased markedly to 48 µg/mL and 64 µg/mL respectively. In contrast, *Kp*_GLPMUT2 exhibited a lower baseline FOS MIC of 6 µg/mL; however, exposure to GLP increased its MIC to 18 µg/mL. Overall, the GLP-evolved mutants displayed substantially higher FOS MICs than the parental strain, particularly under GLP exposure **(Figure 3a)**. All seven of the FOS-evolved mutants exhibited a baseline FOS MIC of >1,024 µg/mL (100 times the MIC of the type strain), which remained >1,024 µg/mL in the presence of a sub-inhibitory concentration GLP **(Figure 3b)**.

Together, these findings indicate that sublethal GLP exposure broadly elevates FOS resistance in *Klebsiella,* with the most pronounced effects observed in the GLP-evolved mutants.

### Comparative analysis of bacterial resistance mechanism to FOS and SNPs identified in *Kp*_FOSMUT3 and *Kp*_GLPMUT2

A comparison of the known bacterial resistance mechanisms against FOS, along with those SNPs identified in *Kp*_FOSMUT3 showed some features of note **(Figure 4).**

**Figure 4.**
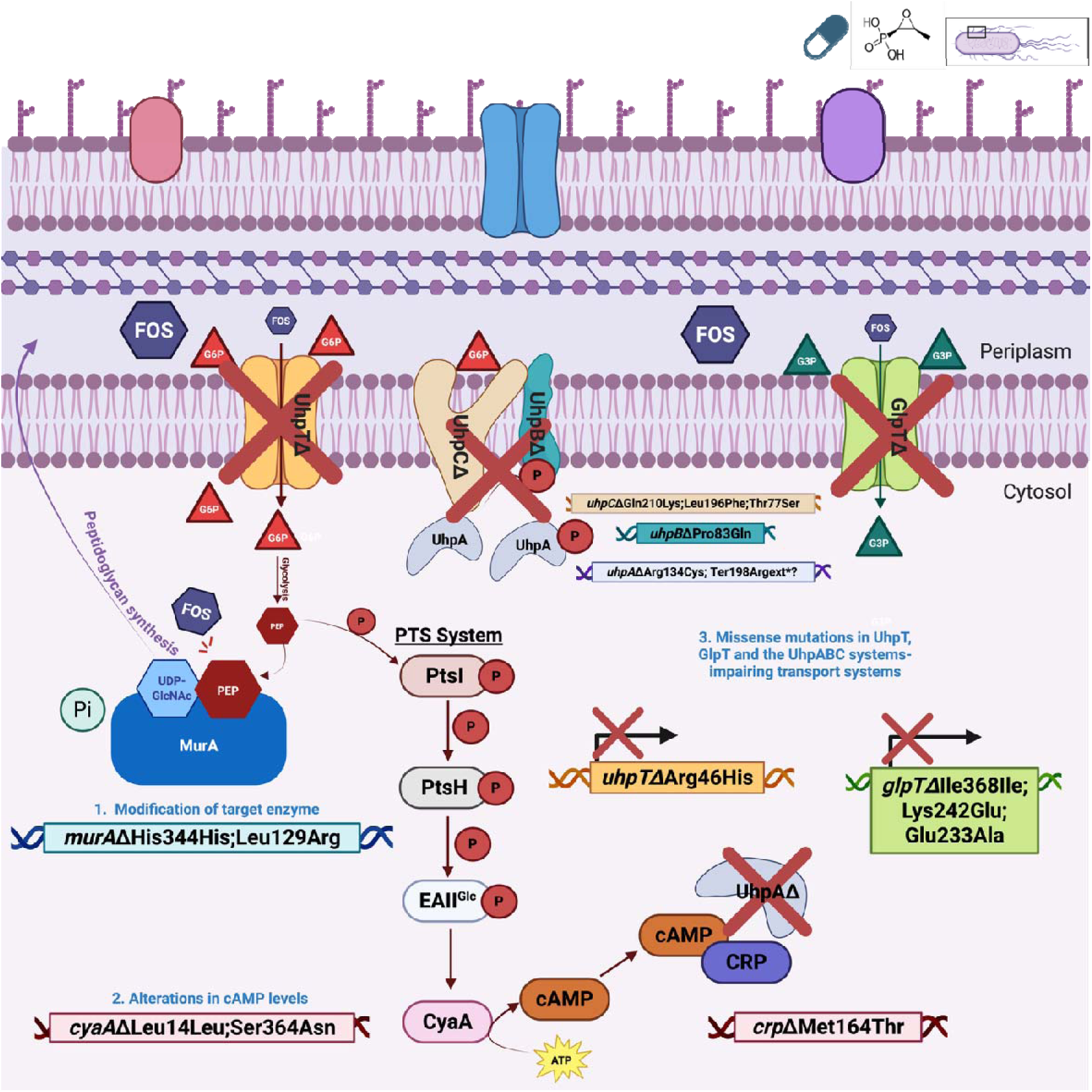
The genetic changes detected in the isogenic FOS-evolved mutant ***Kp*_FOSMUT3** and which by comparison exhibited similar genetic alterations to other *Enterobacteriaceae* that were previously selected on FOS.

Firstly, ***Kp*_FOSMUT3** acquired mutations in the FOS target gene *murA*. This mutant harboured two substitutions, Leu129Arg and the synonymous His344His. While the synonymous change is unlikely to alter protein structure directly, the Leu129Arg substitution may influence MurA conformation or substrate accessibility, potentially reducing FOS binding while preserving the first committed step of peptidoglycan biosynthesis.

Full expression of the sugar-phosphate transporters GlpT and UhpT requires elevated intracellular levels of cAMP, which is synthesised by CyaA and signals through the cAMP receptor protein, CRP (11, 12). In *Kp*_FOSMUT3, two substitutions were identified in *cyaA* (Ser364Asn and the synonymous Leu14Leu), together with a nonsynonymous mutation in *crp* (Met164Thr). Mutations affecting this cAMP-CRP regulatory axis have previously been associated with increased FOS resistance (76), primarily through reduced transcription of *glpT* and *uhpT*. cAMP functions as a global regulatory signal of carbon availability. Disruption of cAMP synthesis or CRP-mediated signalling effectively mimics a carbon-replete state, suppressing the expression of alternative carbon uptake systems, including FOS import pathways, and thereby limiting intracellular drug accumulation (77).

*uhpT* transcription requires UhpA and is positively regulated by cAMP-CRP, which stabilises open promoter complexes and enhances transcription initiation (78). *Kp*_FOSMUT3 carried multiple mutations affecting the *uhp* regulatory and transport system. These included a complex *uhpA* variant (Arg134Cys together with a disruptive stop-loss mutation, Ter198Argext*\**), a nonsynonymous substitution in the sensor kinase *uhpB* (Pro83Gln), three substitutions in receptor-transporter *uhpC* (Thr77Ser, Leu196Phe, and Gln210Lys), and a conserved missense mutation in G6P permease *uhpT* (Arg46His), consistent with strong selective pressure to impair FOS uptake. Inactivation of FOS uptake systems reduces susceptibility, and experimental comparisons indicate that loss of *uhpT* may, in some species (for example *S. aureus*), have a larger effect on FOS resistance than loss of *glpT*, highlighting differential contributions of these transporters to antibiotic entry (79).

The *glpTQ* operon contains multiple cAMP-CRP binding sites and is subject to dual regulation, being repressed by GlpR and activated by cAMP-CRP (80). In *Kp*_FOSMUT3, three SNPs were identified within *glpT* (Ile368Ile, Lys242Glu, and Glu233Ala), indicating potential modulation of the permease. In contrast, in *P. aeruginosa*, where *uhpT* was absent, *glpT* serves as the sole FOS uptake route, and inactivation mutations in *glpT* represent the dominant mechanism of FOS resistance (8).

Taken together, these findings indicate that although individual mutations affecting MurA structure, cAMP-CRP regulation, or FOS transport can each confer partial resistance, their convergence within *Kp*_FOSMUT3 generates a multilayered resistance phenotype involving target modification, impaired antibiotic uptake, and global metabolic reprogramming, culminating in the very high FOS MIC observed (>1,024 µg/mL).

In contrast, comparison of known FOS resistance mechanisms along with a review of the SNPs detected in the GLP-evolved mutant, *Kp*_GLPMUT2, also highlighted overlapping mechanisms **(Figure 5).**

**Figure 5.**
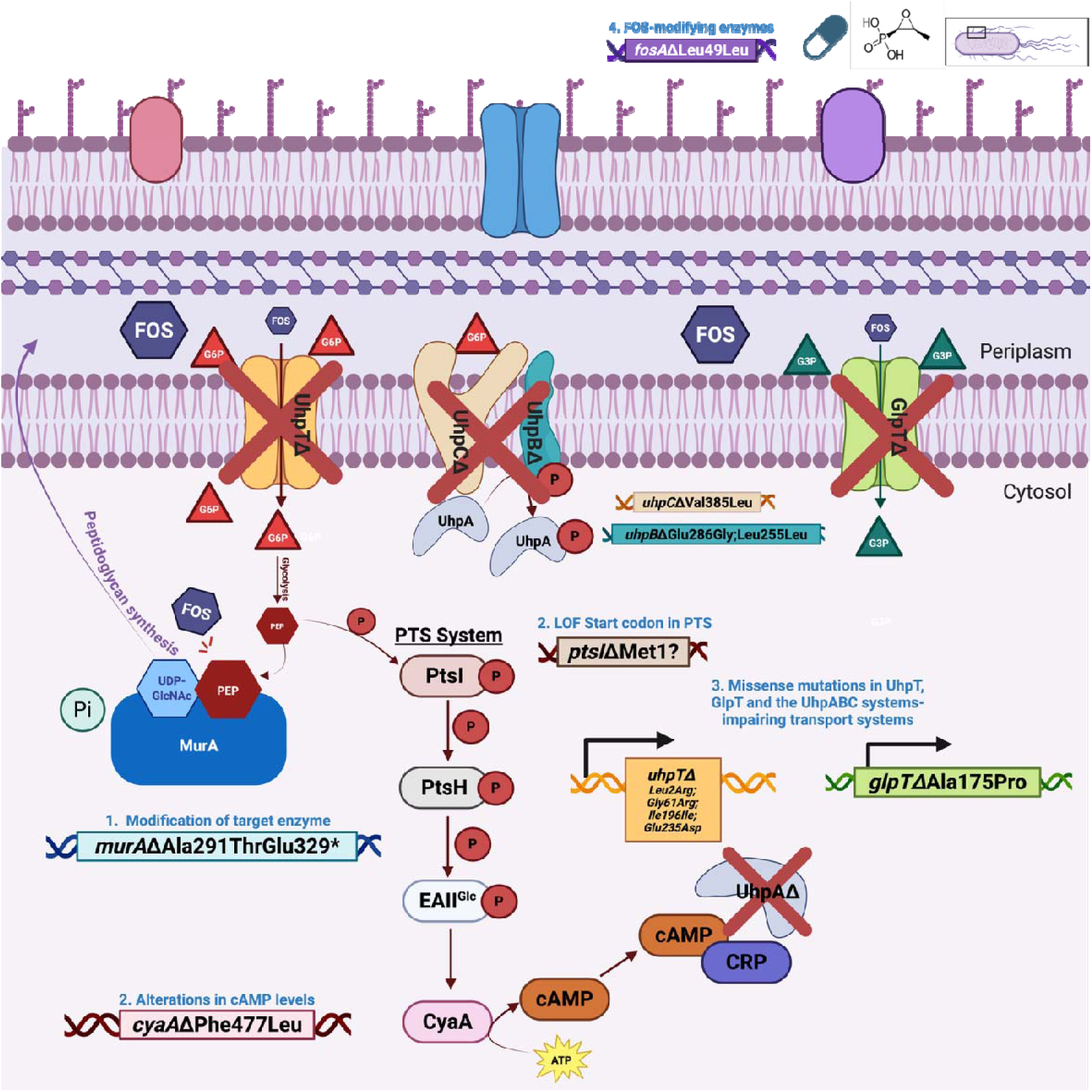
The genetic changes detected in the isogenic GLP-evolved mutant ***Kp*_GLPMUT2** and which by comparison exhibited similar genetic alterations to other *Enterobacteriaceae* that were previously selected on FOS.

***Kp*_GLPMUT2** carried mutations in the FOS target gene *murA*, including a missense substitution (Ala291Thr) and a stop-gained variant (Glu329*). Residue 329 is located within a loop (329–330) that participates directly in interdomain hydrogen bonding with residues 116-119, contributing to stabilisation of the closed conformation of MurA that is required for proper catalytic function and controlled access to the active site. Disruption at this position is therefore expected to impair interdomain communication and alter active-site accessibility. In contrast, residue 291 lies within the C-terminal domain near interdomain contact regions involved in conformational flexibility rather than direct catalytic interactions; substitutions at this site are likely to perturb MurA structural dynamics and stability (17). These *murA* variants in *Kp*_GLPMUT2 are predicted to be substantially more structurally disruptive than the *murA* changes previously observed in *Kp*_FOSMUT3. However, the stop-gained mutation at Glu329 in *murA* is not lethal because this strain underwent a large-scale genomic duplication during the evolution experiment. Gene presence/absence analysis confirmed that *Kp*_GLPMUT2 possessed a second, intact copy of *murA* that maintains essential peptidoglycan synthesis. This duplication likely arose *via* unequal homologous recombination under selective pressure (81), providing the genetic redundancy necessary for the truncated *murA* allele to persist.

In *Kp*_GLPMUT2, a missense substitution was identified in *cyaA* (Phe477Leu), which may influence intracellular cAMP levels, although no mutations were detected in *crp*. Notably, *Kp*_GLPMUT2 also harboured a disruptive mutation affecting the initiator codon (Met1) of *ptsI*, which is predicted to impair or abolish translation of enzyme I, the first and essential component of the PTS system.

The PTS operon initiates carbohydrate uptake through Enzyme I (PtsI)-, and the histidine-containing phosphocarrier protein HPr (PtsH)-mediated phosphorylation before diverging into the sugar-specific enzyme II complexes, including those for glucose, mannitol, mannose, and lactose/chitobiose (82, 83). Inactivation of *ptsI* abolishes EI activity, thereby disabling the PTS and preventing utilisation of PTS-dependent sugars, often resulting in broad defects in growth on multiple carbon sources and altered global regulatory responses. For example, deletion of *ptsI* in group A streptococci renders them unable to grow on both PTS (such as galactose, trehalose, and fructose) and certain non-PTS carbon sources [such as glycerol, G6P and fructose-6-phosphate (F6P)], demonstrating the system’s essential role in carbon metabolism (84). Given the close integration of the PTS with global metabolic regulation, including catabolite control and cAMP signalling (85), disruption of *ptsI* is likely to have pleiotropic effects on carbon utilisation, energy balance and regulatory networks, with potential downstream consequences for transporter expression and FOS susceptibility in *Kp*_GLPMUT2.

Further, *Kp*_GLPMUT2 harboured multiple mutations affecting the *uhp* regulatory and transport system. No variants were detected in *uhpA*; however, *uhpB* carried one nonsynonymous and one synonymous substitution (Glu286Gly; Leu255Leu), *uhpC* contained a missense mutation (Val385Leu), and *uhpT*, encoding the G6P permease, accumulated three substitutions (Leu2Arg, Gly61Arg, Ile196Ile, and Glu235Asp). The preponderance of mutations recorded within the *uhp* sensing and transport components is consistent with strong selective pressure to reduce G6P-mediated FOS uptake, as in *Kp*_FOSMUT3.

In *Kp_*GLPMUT2, only a single nonsynonymous mutation was detected in *glpT* (Ala175Pro), suggesting limited modulation of the G3P permease. This contrasts with *Kp*_FOSMUT3, in which multiple mutations affecting this FOS uptake pathway were observed.

As described above, CARD analysis indicated that *Kp*_type and all GLP- and FOS-evolved mutants carried the *fosA6* gene. A single SNP was identified in KPN_04759, annotated by NCBI as a *FosA5 family fosfomycin resistance glutathione transferase* and by Snippy as a *glyoxalase/bleomycin resistance protein/dioxygenase*, highlighting annotation inconsistencies between databases for *fosA*-like genes. This mutation was synonymous (Leu49Leu) and is therefore unlikely to directly affect protein function.

### Detailed analysis of bacterial resistance mechanism to GLP and SNPs identified in *Kp*_FOSMUT3 and *Kp*_GLPMUT2

Bacterial resistance to GLP arises through multiple mechanisms, including altered EPSPS activity or expression, GLP degradation or detoxification, and reduced uptake or enhanced efflux (23). One mechanism by which bacteria can acquire GLP resistance is through mutations in the EPSPS active site that reduce or prevent GLP binding **(Figure 6).**

**Figure 6.**
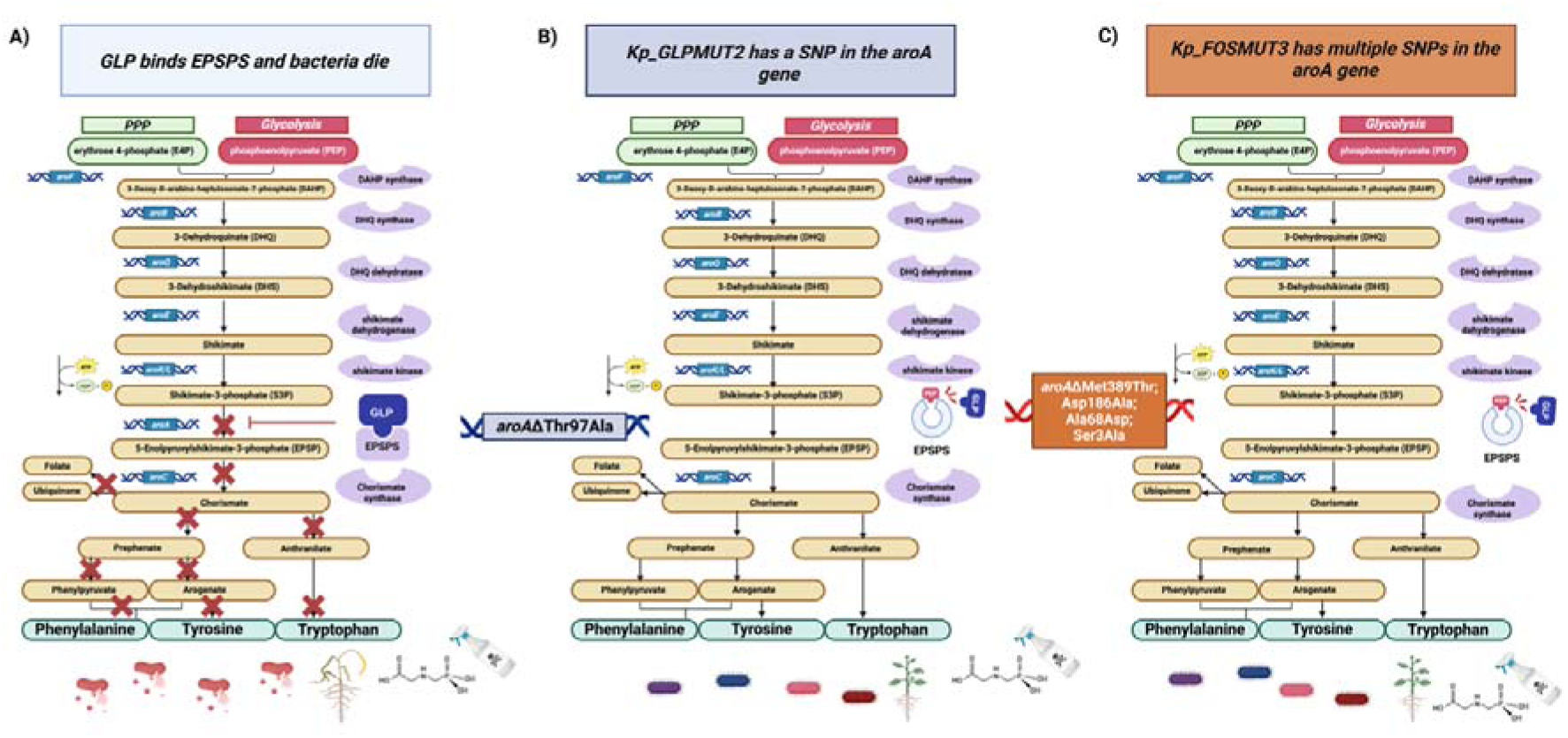
Altered EPSP synthase activity. **A**) A disrupted shikimate pathway by GLP results in plant and bacterial death. GLP mimics PEP and binds the active site of EPSPS with a positive charge; **B**) The *Kp*_GLPMUT2 has an alanine substitution at position 97 in the *aroA* gene; **C**) The *Kp*_FOSMUT3 has multiple substitutions in the *aroA* gene.

*Kp*_FOSMUT3 carried four SNPs in the *aroA* gene encoding EPSPS; Ser3Ala; Ala68Asp; Asp186Ala; Met389Thr. Classification using the EPSPS Class analysis tool (http://ppuigbo.me/programs/EPSPSClass/) identified the EPSPS sequence of *Kp*_FOSMUT3 as Class I (alpha), the same class as the parental *Kp*_type, whereas *Kp*_GLPMUT2 (Thr97Ala) was classified as Class I (beta). Both subclasses belong to the Class I EPSPS group, and which are generally considered to be susceptible to GLP. Although none of the identified SNPs in *Kp*_FOSMUT3 occur within amino acids 90-104, which form a highly conserved motif in the EPSPS active site, the presence of four substitutions within *aroA* is nevertheless notable. Previous work attempting to clone a GLP-resistant *aroA* Cys358Ala allele from *E. coli* demonstrated extensive genetic instability, with most clones acquiring additional missense mutations, frameshifts, or C-terminal extensions. Complementation analyses further showed that an AroA variant carrying multiple substitutions (Gly14Val; Arg120Ser; Asp313Gly; Thr381Ala; Met389Thr; Thr409Ser) was catalytically inactive (86). One of the acquired mutations targeted the aspartate residue at position 313, which has been shown to play a key role in EPSPS catalytic activity (87).

The Thr97Ala SNP in *Kp*_GLPMUT2 is supported by two lines of evidence: (88) showed that disrupting Thr97 in the Thr97Ile/Pro101Ser TIPS double mutation breaks a conserved hydrogen bond that reshapes the EPSPS active site and alters GLP and PEP interactions, while (89) demonstrated that adding a methyl group from Ala at the adjacent Gly96 position sterically impairs GLP binding. By analogy with these characterised mutants, the Thr97Ala substitution in *Kp*_GLPMUT2 could, in principle, destabilise local active-site structure and reduce GLP affinity.

Interestingly, there appears to be an inverse relationship in target-site adaptation, as *Kp*_GLPMUT2 evolved potentially disruptive mutations in MurA **(Figure 5)** while *Kp*_FOSMUT3 evolved potentially disruptive mutations in EPSPS **(Figure 6)**. Nonetheless, *Kp*_FOSMUT3 remains viable and metabolically competent. Compared with the previously described inactive variant by Eschenburg and colleagues, *Kp*_FOSMUT3 harboured fewer SNPs and when they arose they avoided critical catalytic residues. Instead, these substitutions may subtly alter enzyme structure or dynamics, potentially contributing to reduced GLP susceptibility while preserving essential catalytic function.

A second mechanism by which bacteria can acquire GLP resistance involves metabolising or detoxifying GLP and utilising it as a phosphorus source, either *via* the C-P lyase pathway or through an oxygen-dependent oxidative cleavage reaction **(Figure 7).**

**Figure 7.**
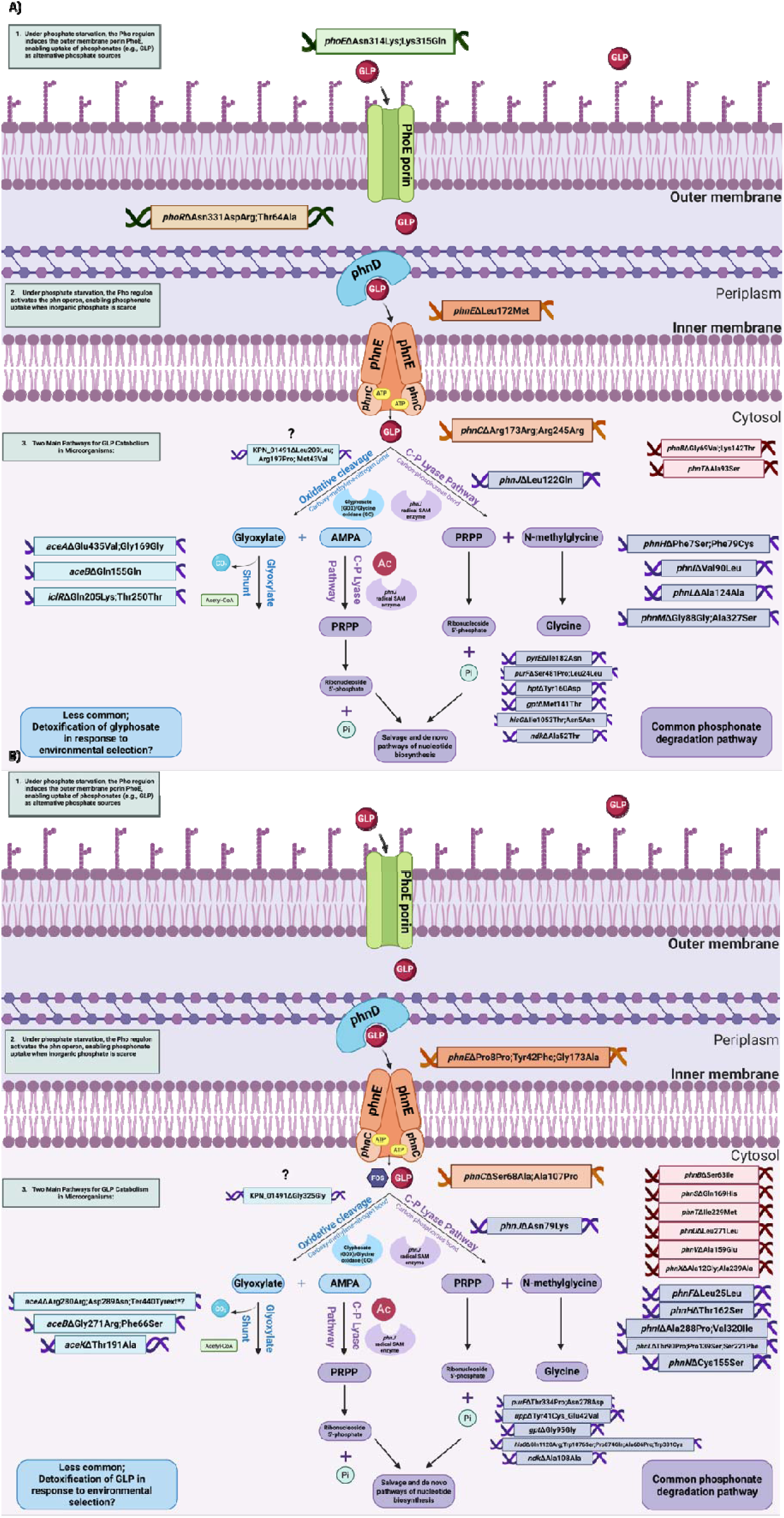
Metabolic strategies in Gram-negative bacteria responsible for GLP degradation and which may serve as a resistance mechanism, particularly under phosphate-limited conditions. **A)** The *Kp*_GLPMUT2 has SNPs throughout the entire degradation pathway, **B)** as does *Kp*_FOSMUT3.

A clear difference noted between *Kp*_GLPMUT2 and *Kp*_FOSMUT3 is the absence of SNPs in PhoE and PhoBR in the FOS mutant, implying that modulation of membrane-associated GLP or phosphorus transport is not selectively favoured in the FOS mutant. Under phosphate-limiting conditions, the sensor histidine kinase PhoR autophosphorylates and transfers a phosphate to the response regulator PhoB, which then binds the Pho box and activates transcription of Pho regulon operons (90). In *Kp*_GLPMUT2, two missense substitutions were identified in PhoR: Asn331Asp and Thr64Ala. PhoR is a 431-AA protein, and residue Asn331 lies within the C-terminal catalytic ATP-binding domain (residues 268-431), which mediates PhoR autophosphorylation under low-phosphate conditions. In contrast, Thr64 is positioned between the N-terminal transmembrane domain (residues 10-59) and the adjacent cytoplasmic charged region (residues 70-100), suggesting it may influence signal transduction rather than catalytic activity (91). Activation of the Pho regulon also induces outer membrane proteins such as PhoE, which facilitate the uptake of phosphorus containing compounds into the periplasm (90).

The *phn* operon encodes the C-P lyase system and associated transporter components. The ABC transporter comprises PhnD (periplasmic binding protein), PhnE (membrane spanning protein), and PhnC (ATP binding protein), which together mediate phosphonate uptake, including GLP. The downstream PhnGHIJK proteins form the core of phosphonate degradation: PhnGHI act as components of purine ribonucleoside triphosphate phosphonylase; PhnJ (EC 4.7.1.1) performs the key C-P bond-cleavage reaction; PhnI contributes catalytically to the PhnGHJK complex; PhnL likely functions as a nucleotide binding protein; and PhnM acts as a triphosphoribosyl diphosphohydrolase that generates Phosphoribosyl pyrophosphate (PRPP) in cooperation with PhnN (92). Consistent with the presence of a C-P bond in FOS, *Kp*_FOSMUT3 exhibited a substantial enrichment of SNPs in the C-P lyase pathway, including the *phnCDEFGHIJKLMNOP* operon (purple colours in **Figure 7B**) and other associated *phn* genes (*phnBSTUVX*; red colours in **Figure 7B**). In *Kp*_FOSMUT3 missense mutations were identified in *phnE* (Tyr42Phe;Gly173Ala), *phnC* (Ser68Ala;Ala107Pro), *phnH* (Thr162Ser), *phnI* (Ala288Pro;Val320Ile), *phnJ* (Asn79Lys), *phnL* (Thr90Pro;Pro139Ser;Ser221Phe), and *phnN* (Cys155Ser)), with several synonymous substitutions also recognised (*phnE*: Pro8Pro, *phnF*: Leu25Leu). *Kp*_FOSMUT3 exhibited additional SNPs in *phnF* and *phnN* and no longer carried the *phnM* SNP present in *Kp*_GLPMUT2. Transposon mutagenesis has shown that insertions in *phnC*, *phnD*, *phnE*, *phnG*, *phnH*, *phnI*, *phnJ*, *phnK*, *phnL*, *phnM*, or *phnP* abolish phosphonate utilisation, whereas insertions in *phnF*, *phnN*, or *phnO* do not impair growth on phosphonates (93).

Indeed, novel mechanisms underlying FOS resistance were identified for phosphonate catabolism (the C-P lyase *phnC-M* operon), and the phosphate transporter PstSACB (94), thereby expanding known FOS resistance mechanisms and further linking FOS and GLP resistance. Furthermore, *Kp*_FOSMUT3 harboured SNPs in *pstSACB* (*pstS*: Thr171Asn, Asp138His, Ser64Cys; *pstA*: Leu267Gln; *pstC*: Ala27Gly), suggesting reliance on an alternative inorganic phosphate importer in the absence of PhoE. In contrast, *Kp*_GLPMUT2 carried SNPs only in *pstS* (Thr6Ser, Leu266Leu, Pro276Pro).

*Kp*_FOSMUT3 contained a missense variant in *phnB* (Ser63Ile), which is identified as *Glyoxalase/fosfomycin resistance/dioxygenase domain-containing protein* in the MGH 78578 (*Kp*_type) reference used for Snippy analysis and annotated as *VOC family metalloprotein YjdN* in the NCBI database. The vicinal oxygen chelate (VOC) superfamily consists of structurally related enzymes characterised by paired βαβββ motifs that create a metal-binding site with two or three accessible coordination positions. This architecture enables direct participation of the metal ion in catalysis, supporting a wide range of reactions, including isomerisation (glyoxalase I), epimerisation (methylmalonyl-CoA epimerase), oxidative C-C bond cleavage (extradiol dioxygenases), sequestration (bleomycin resistance proteins) and nucleophilic substitution (FOS resistance proteins) (95). *Kp*_GLPMUT2 carried SNPs also in *phnB* (Gly69Val;Lys142Thr). Together, SNPs in *phnB* in both *Kp*_FOSMUT3 and *Kp*_GLPMUT2 implicate VOC-family metalloproteins at the intersection of phosphonate metabolism and resistance, highlighting a shared metal-dependent catalytic framework that links phosphonate catabolism (*phn* genes) with FOS resistance (FOS resistance proteins), and GLP and FOS detoxification mechanisms.

Further*, Kp*_FOSMUT3 harboured multiple SNPs in the *phnSTUV* operon, (phosphonatase pathway) which encodes the 2-aminoethylphosphonate (2AEP) transporter, including variants in the periplasmic binding protein *phnS* (Gln169His), the ATPase *phnT* (Ile229Met), and the membrane components *phnU* (Leu271Leu) and *phnV* (Ala159Glu). This transporter enables uptake of 2AEP for use as alternative carbon, nitrogen and phosphorus sources, independently of inorganic phosphate availability (96). In addition, *Kp*_FOSMUT3 carried SNPs in *phnX*, encoding phosphonoacetaldehyde hydrolase (EC 3.11.1.1) (Ala12Gly; Ala239Ala), which catalyses C-P bond cleavage during 2AEP catabolism (97). In contrast, *Kp*_GLPMUT2 contained a SNP only in *phnT* (Ala93Ser), suggesting that acquisition of alternative phosphorus from environmental phosphonate sources, such as 2AEP, is of greater importance in the FOS-selected background.

Considering the PRPP-dependent phosphoribosyltransferase (PRTase) enzymes that function across both salvage and *de novo* biosynthetic pathways for purine, pyrimidine, and pyridine nucleotides, as well as histidine (98), *Kp*_FOSMUT3 also carried multiple SNPs in genes associated with these metabolic routes. As noted earlier, *phnN*, which generates PRPP, contained a Cys155Ser variation. In *de novo* nucleotide biosynthesis, PRPP serves as a central precursor. In pyrimidine biosynthesis, PRPP reacts with orotate through the action of *pyrE* (EC 2.4.2.10) (which was devoid of any SNP). In purine biosynthesis, the pathway is initiated when PRPP reacts with ammonia derived from glutamine *via purF* (EC 2.4.2.14) (Thr334Pro;Asn278Asp), ultimately producing inosine 5′-monophosphate (IMP) through a 10-step process.

In salvage pathways, preformed nucleobases are converted to nucleotides by reaction with PRPP. Uracil salvage occurs *via* the uracil PRTase *upp* (EC 2.4.2.9) (complex missense variant Tyr41Cys_Glu42Val), which attaches the N-1 atom of uracil to the C-1 position of PRPP. Three purine PRTases also function in salvage: *apt* (EC 2.4.2.7) (adenine-specific; no SNP), *hpt* (EC 2.4.2.8) (prefers hypoxanthine and guanine; no SNP), and *gpt* (EC 2.4.2.22) (acts on xanthine and guanine; Gly95Gly). Histidine biosynthesis also begins with PRPP, through the ATP PRTase *hisG* (EC 2.4.2.17) which had the following SNPs (Trp381Cys;Ala606Pro;Pro874Gln;Trp1076Ser;Gln1120Arg), although this pathway is inactive when histidine is available (99). Conversion of nucleoside diphosphates to triphosphates is catalysed by *ndk* (EC 2.7.4.6) (AA substitution identified, Ala108Ala), which maintains balanced nucleoside triphosphate pools.

In summary, both *Kp*_FOSMUT3 and *Kp*_GLPMUT2 accumulated SNPs in PRPP-dependent PRTase pathways, underscoring nucleotide metabolism as a shared adaptive target under FOS and GLP selection. However, *Kp*_FOSMUT3 showed broader and more functionally consequential variation across PRPP generation (*phnN*), *de novo* purine synthesis (*purF*; purine only), and uracil nucleotide salvage (*upp*), consistent with a more extensive rewiring of PRPP flux and downstream biosynthetic demands. In contrast, *Kp_*GLPMUT2 displayed a more limited and pathway-specific pattern, with mutations concentrated in *de novo* nucleotide synthesis (*pyrE*, *purF*), purine salvage (*hpt*, *gpt*), and *ndk*, while retaining an unaltered PRPP supply *via phnN*. However, both strains converged on multiple SNPs in *hisG*, indicating histidine biosynthesis as a shared target of adaptation. Together, these patterns suggest that FOS selection imposes stronger pressure on PRPP availability and utilisation across interconnected anabolic and salvage pathways, whereas GLP selection favours more localised modulation of nucleotide metabolism, reflecting distinct metabolic constraints underpinning resistance to these compounds.

Oxidative cleavage of the C-N bond by the FAD-dependent oxidoreductases glycine oxidase (GO; EC 1.4.3.19) and glyphosate oxidase (GOX; EC 1.5.3.23), is well characterised, particularly in Gram-positive bacteria (100, 101). To determine whether *Klebsiella* might encode similar enzymes capable of GLP degradation, the predicted amino acid sequences of *Kp*_type, *Kp*_GLPMUT2, and *Kp*_FOSMUT3 were compared by BLAST with known GLP-degrading oxidases (GOX, GenBank ACZ58378.1; GO, GenBank AEW04802.1; *goW*, GenBank OM867748). No significant similarity was detected, indicating that *Klebsiella* does not appear to possess canonical glycine nor GLP oxidases.

In contrast, *Enterobacteriaceae* primarily utilise the glycine cleavage system, encoded by the *gcvTHP* operon and regulated by cAMP-CRP (102), rather than FAD-dependent glycine oxidases. An alternative GLP detoxification strategy has been described in plants: *E. colona* evolved GLP resistance through expression of an aldo-keto reductase (AKR) capable of metabolising GLP. Functional assays demonstrated that EcAKR4-1 converts GLP into AMPA and glyoxylate when expressed in *E. coli* (103). To assess whether *Klebsiella* might encode a related enzyme, the amino acid sequences of *Kp*_type, *Kp*_GLPMUT2, and *Kp*_FOSMUT3 were compared with the plant AKR enzymes (EcAKR4-1, GenBank MK592097; EcAKR4-2, GenBank MK592098). No significant similarities were found.

However, several SNPs were identified in a predicted aldo/keto reductase oxidoreductase gene, KPN_01491 in *Kp*_GLPMUT2 (Leu209Leu; Arg197Pro; Met43Val) and *Kp*_FOSMUT3 (Gly325Gly) **(Figure 7)**. Although the functional implications of these mutations are unclear at this time, AKR enzymes are known to participate broadly in xenobiotic detoxification pathways. Thus, this locus represents a potentially important candidate for further investigation into GLP and FOS metabolism or tolerance in *Klebsiella*.

Oxidative cleavage of GLP produces glyoxylate and AMPA, and the former can subsequently enter the glyoxylate shunt. The glyoxylate shunt enables cells to utilise acetate and fatty acids for gluconeogenesis, and is defined by two key enzymes: isocitrate lyase (ICL; EC 4.1.3.1), encoded by *aceA*, which cleaves isocitrate into succinate and glyoxylate, and malate synthase (MS; EC 2.3.3.9), encoded by *aceB*, which condenses glyoxylate with acetyl-CoA to form malate. The pathway is negatively regulated by IclR, a transcriptional repressor of *aceA* and *aceB*. IclR activity is controlled by metabolic signals: glyoxylate and PEP stabilise its inactive dimeric form, whereas pyruvate favours the active tetrameric form; additionally, PEP acts as an uncompetitive inhibitor of ICL activity (104, 105).

*Kp*_FOSMUT3 carried multiple variants in *aceA*, including loss of the native stop codon resulting in C-terminal extension (Arg280Arg, Asp289Asn, Ter440Tyrext*), as well as SNPs in *aceB* (Gly271Arg, Phe66Ser). In contrast to *Kp*_GLPMUT2, *Kp*_FOSMUT3 additionally harboured a SNP in *aceK* (Thr191Ala) and lacked mutations in the negative regulator *iclR*. As *aceK* encodes the bifunctional isocitrate dehydrogenase kinase/phosphatase that controls carbon flux between the TCA cycle and the glyoxylate shunt, these mutations suggest altered regulation of isocitrate partitioning (106). Interestingly, deletion of *aceA* in *P. aeruginosa* improved survival under oxidative and antibiotic stress, suggesting that downregulation of this pathway can be advantageous (107). Collectively, both *Kp*_FOSMUT3 and *Kp*_GLPMUT2 appear to downregulate glyoxylate shunt activity, albeit through distinct genetic mechanisms.

A third bacterial mechanism of GLP resistance involves reduced GLP import and enhanced export **(Figure 8)**. In this context, the SNPs identified in *Kp*_FOSMUT3 were therefore considered in relation to potential alterations in transport and efflux systems.

**Figure 8.**
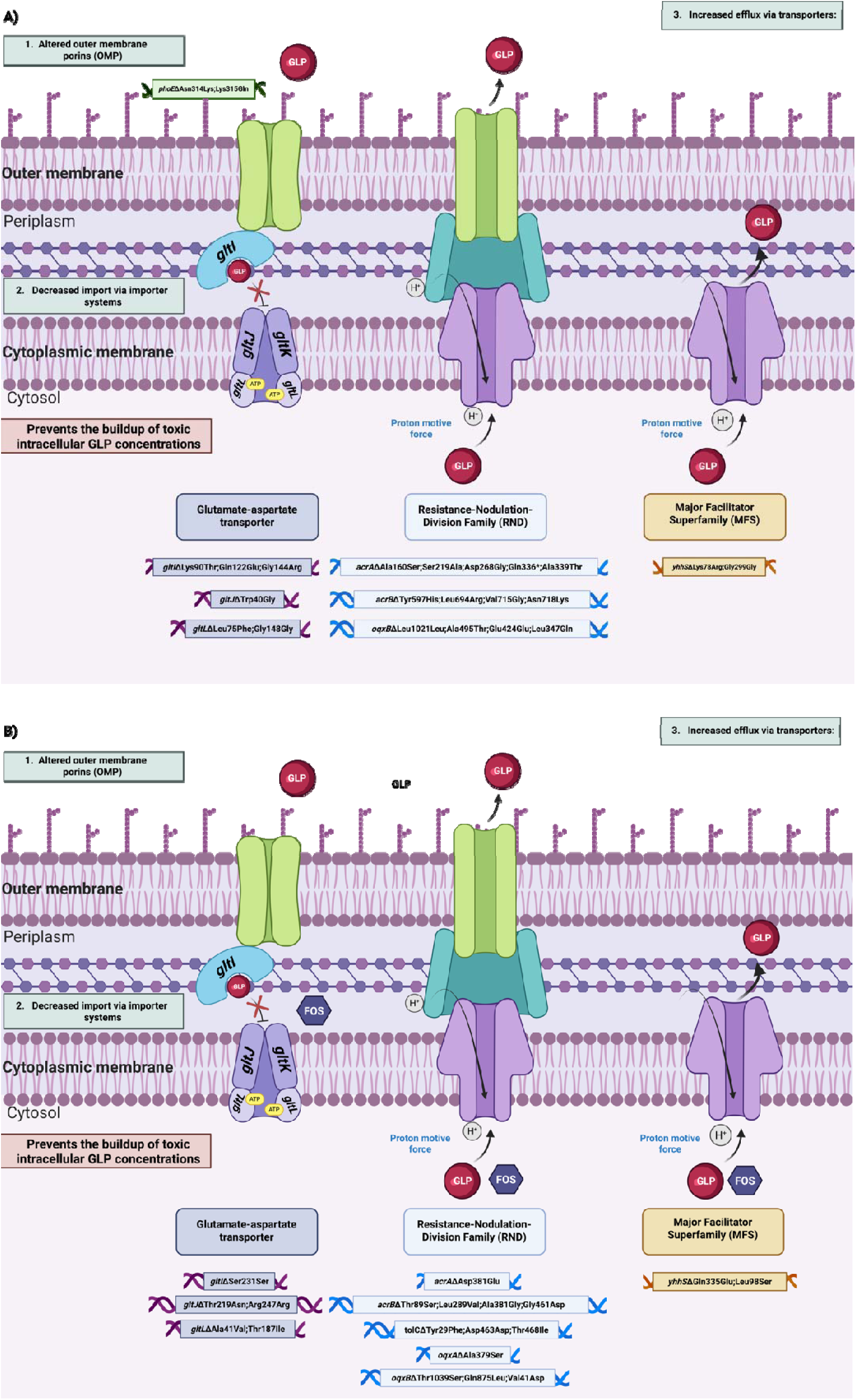
Bacteria can evade GLP toxicity through modifications in membrane transport systems that limit intracellular accumulation of the compound. **A)** The *Kp*_GLPMUT2 has SNPs in many of the transporters, **B)** as does *Kp*_FOSMUT3.

*Loss-of-function* mutations in glutamate transporters are known to confer resistance to GLP (108). *Kp*_FOSMUT3 harboured SNPs in multiple components of the glutamate/aspartate transport system, including the periplasmic binding protein *gltI* (Ser231Ser), the membrane-spanning protein *gltJ* (Thr219Asn;Arg247Arg), and the ATP-binding protein *gltL* (Ala41Val;Thr187Ile). This suggests that both mutants may limit GLP and FOS uptake by disabling glutamate transport while concurrently limiting intracellular glutamate availability, potentially shifting reliance toward phosphonate import and subsequent breakdown as an alternative phosphorus source.

The most studied and predominant efflux pumps in *Klebsiella* are those of the Resistance-Nodulation-Division Family (RND)-type efflux systems including AcrAB-TolC and OqxAB (109). *Kp*_FOSMUT3 accumulated additional SNPs in AcrAB-TolC and OqxAB. Variants were identified in *acrA* (Asp381Glu), *acrB* (Thr89Ser, Leu289Val, Ala381Gly, Gly461Asp), *tolC* (Tyr29Phe, Asp463Asp, Thr468Ile), and the AcrAB repressor *acrR* (Glu61Asp). Mutations were also identified in the *oqxAB* efflux system, including *oqxA* (Ala379Ser) and *oqxB* (Thr1039Ser, Gln875Leu, Val41Asp). In contrast, *Kp_*GLPMUT2 lacked SNPs in *tolC*, *acrR*, and *oqxA*, suggesting that FOS resistance more strongly relies on enhanced RND-type efflux activity. Furthermore, exposure to GLP has been shown to induce SoxS-mediated overexpression of the AcrAB-TolC efflux system in *E. coli* (110).

Overexpression of the membrane transporter *yhhS* in *E. coli* and *Pseudomonas* species has been shown to confer high-level resistance to GLP (111). *Kp*_FOSMUT3 carried two variants in the GLP-associated MFS exporter *yhhS* (Gln335Glu, Leu98Ser), as did *Kp_*GLPMUT2 (Gly299Gly;Lys78Arg), strongly suggesting overlapping export mechanisms underlying GLP and FOS resistance.

## Conclusions

Data obtained from WGS-based analysis of FOS- and GLP-evolved mutants, suggested that exposure to GLP and FOS selects for interconnected adaptive pathways that promote both increased AMR and broad physiological reprogramming in *Klebsiella*. Although the MIC for GLP-evolved mutants peaked at only double that of the *Kp_type* strain, FOS-evolved mutants exhibited a 100-fold increase. This disparity suggests a distinct metabolic bottleneck inherent to GLP resistance that is notably absent in the FOS evolutionary pathway. Furthermore, while alternative resistance mechanisms that remain to be elucidated exist in non-hypermutated lineages, the acquisition of potentially lethal mutations in primary targets necessitates a hypermutator phenotype to facilitate the compensatory genomic shifts required for viability. Sub-lethal GLP exposure consistently elevated FOS resistance across the study collection, with the most pronounced effects observed in the GLP-evolved mutants.

The resistance profiles observed between the evolved lineages underscore two distinct evolutionary trajectories toward FOS tolerance. In *Kp*_FOSMUT3, the convergence of mutations affecting the MurA target, global carbon regulation (cAMP-CRP), and FOS transport produced a multilayered resistance phenotype. This combination of target modification, alterations in transport mechanisms, and extensive metabolic reprogramming culminated in high-level resistance MIC_FOS_ (>1,024 µg/mL).

Conversely, the trajectory of *Kp*_GLPMUT2 was dictated by its hypermutator status, driven by high-impact truncations in *dnaQ* and *mutS*. This elevated mutation rate facilitated a rare gene duplication event, providing a second, functional copy of the essential *murA* gene important for FOS resistance. This genetic redundancy permitted the acquisition of a normally lethal Glu329 stop-gain* in the primary *murA* allele, alongside mutations in central carbon regulation (*cyaA* and *ptsI*) and the *uhp* transport system. While these adaptations impose significant metabolic constraints, evidenced by a modest baseline MIC_FOS_ (6 µg/mL), the strain remains highly responsive to environmental cues. The observed shift to an MIC_FOS_ of 18 µg/mL under sub-lethal GLP exposure suggests a ’priming’ effect, where GLP-induced physiological stress transiently enhances the underlying genetic resistance, which was also observed in *Kp*_GLPMUT1 MIC_FOS_ (48 µg/mL) and *Kp*_GLPMUT3 MIC_FOS_ (64 µg/mL). Ultimately, these findings demonstrate that the observed SNPs allow *K. pneumoniae* to navigate intense agrichemical selective pressures.

Comparative analysis of GLP and FOS evolutionary mutants identified a small set of genes repeatedly targeted by selection, indicating convergent adaptation to GLP and FOS stress. Interestingly, many of these SNPs arose independently at the same genomic loci across all mutants, suggestive of a convergent evolution. Shared mutations in *yddH*, *parE*, and the large BapA-like protein (KPN_00994) suggest selection on redox balance, DNA topology-associated fitness, and surface or biofilm-associated traits, rather than direct antibiotic target modification alone. Recurrent alterations in ethanolamine utilisation (*eutA*), citrate metabolism (*citG*), and phosphorylated substrate transport (*uhpT*/*uhpB*) further indicate a coordinated metabolic shift away from host-derived phosphorylated nutrients toward alternative carbon sources such as glycerol. Together with mutations affecting envelope-associated signalling (*yfiR/pal*) and cyclic-di-GMP-regulated adhesion pathways, these findings support a model in which resistance emerges through integrated metabolic rewiring, redox adaptation, and surface remodelling. GLP and FOS resistance in *Klebsiella* appears to be driven by multifactorial adaptive responses that couple altered metabolism and stress tolerance with AMR.

When considering bacterial resistance mechanisms to GLP, the SNP profile of *Kp*_FOSMUT3 supports this multifactorial model of resistance driven by extensive metabolic rewiring. In contrast to the GLP mutant, *Kp*_GLPMUT2, which exhibits modest EPSPS modification alongside Pho-regulated phosphonate uptake and degradation, *Kp*_FOSMUT3 shows broader adaptation of phosphonate catabolism, phosphate acquisition, and PRPP-dependent nucleotide metabolism. These changes are accompanied by altered regulation of the glyoxylate shunt, and enhanced efflux *via* RND-type pumps and the GLP-associated MFS exporter *yhhS*. Furthermore, FOS resistance in *Klebsiella* appears to rely more heavily on enhanced phosphorus scavenging and shared export mechanisms, highlighting overlapping yet distinct metabolic strategies underpinning resistance to FOS and GLP.

Limitations of this work was that the FOS mutant’s MICs exceeded the highest concentrations available on the FOS e-Strips, limiting the complete assessment of MICs in the presence of sub-lethal GLP. Future work includes additional functional characterisation of the identified SNPs to establish definitive genotype-phenotype relationships

Finally, these findings support the conclusion that GLP could act as a potent metabolic co-stressor, and as an antimicrobial-like agent, driving integrated metabolic, redox, and surface adaptations that compromise the efficacy of antimicrobials such as FOS. This work reveals a previously unrecognised, metabolism-based dimension of AMR in *Klebsiella*, with important implications for environmental chemical exposure and AMR evolution.

## Acknowledgements

The generative AI tool ChatGPT (*version GPT-5.2*), developed by OpenAI, was used to assist with language refinement of the manuscript.

I acknowledge BioRender for the production of the figures in this manuscript :

Graphical abstract: Created in BioRender. Wall, K. (2026) https://BioRender.com/utsjv1m

## Author contribution statement

Conceptualisation: KW and SF

Data curation: KW

Formal analysis: KW

Funding acquisition: KW

Investigation: KW, AC, and AK

Methodology: KW and SF

Project administration: KW and SF

Resources: SF

Software: KW and MM

Supervision: SF (KW and AC) and HM (AC)

Validation: KW and AC

Visualisation: KW

Writing: KW

Writing – review and editing: CD, MM, and SF

## Data availability statement

The genome sequences generated in this study are available in the NCBI BioProject database under accession number **PRJNA1423036**. Associated BioSample and Sequence Read Archive (SRA) accession numbers are provided within the BioProject record. To be released upon publication.

