## Supplemental files for "Glyphosate, a herbicide, and fosfomycin, an antibiotic in clinical use- evidence of common selectable genotypes"

**Supplementary files-**

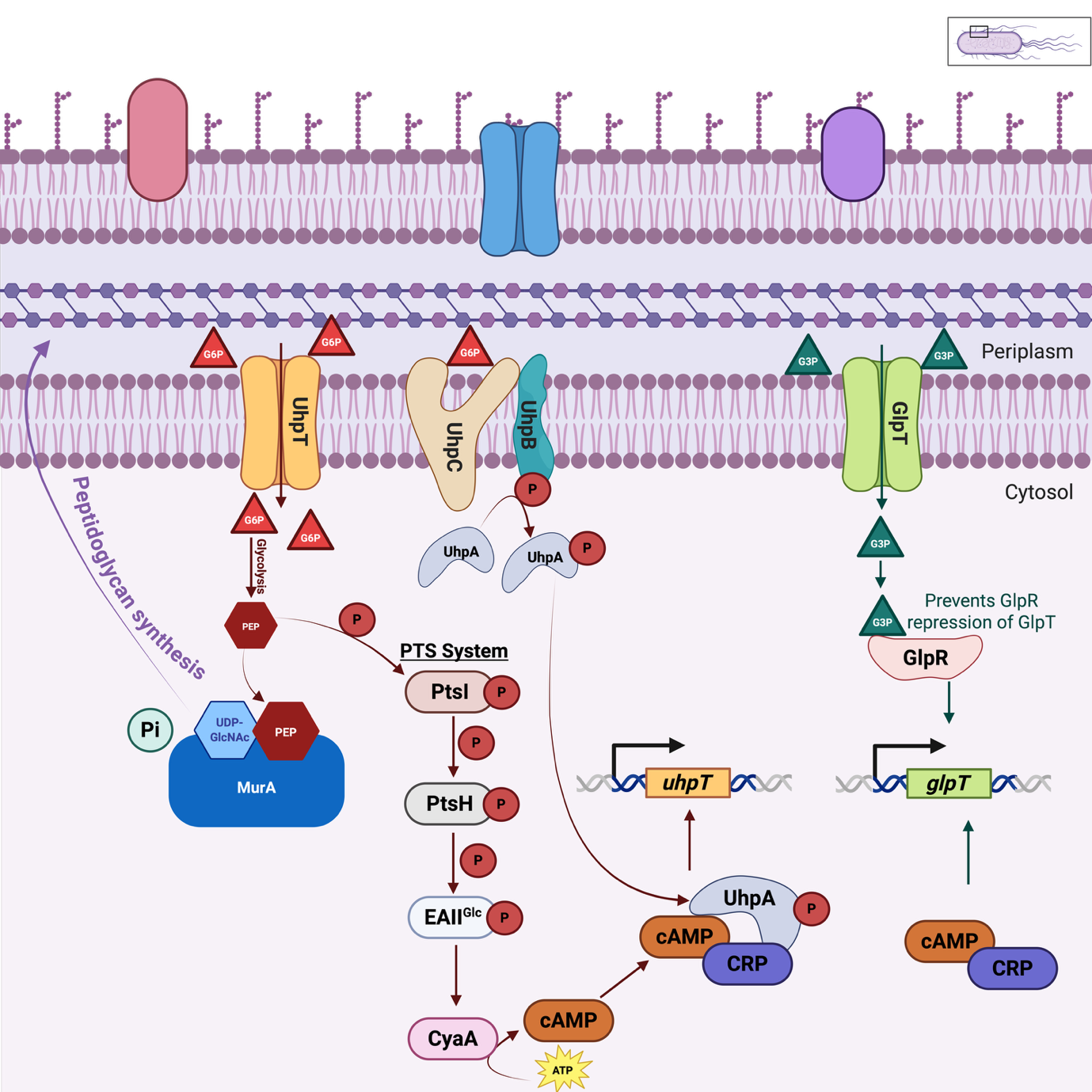

**Figure S1-:** Schematic illustration of the metabolic pathways linking sugar phosphate transporters (UhpT and GlpT) to peptidoglycan biosynthesis in bacteria.

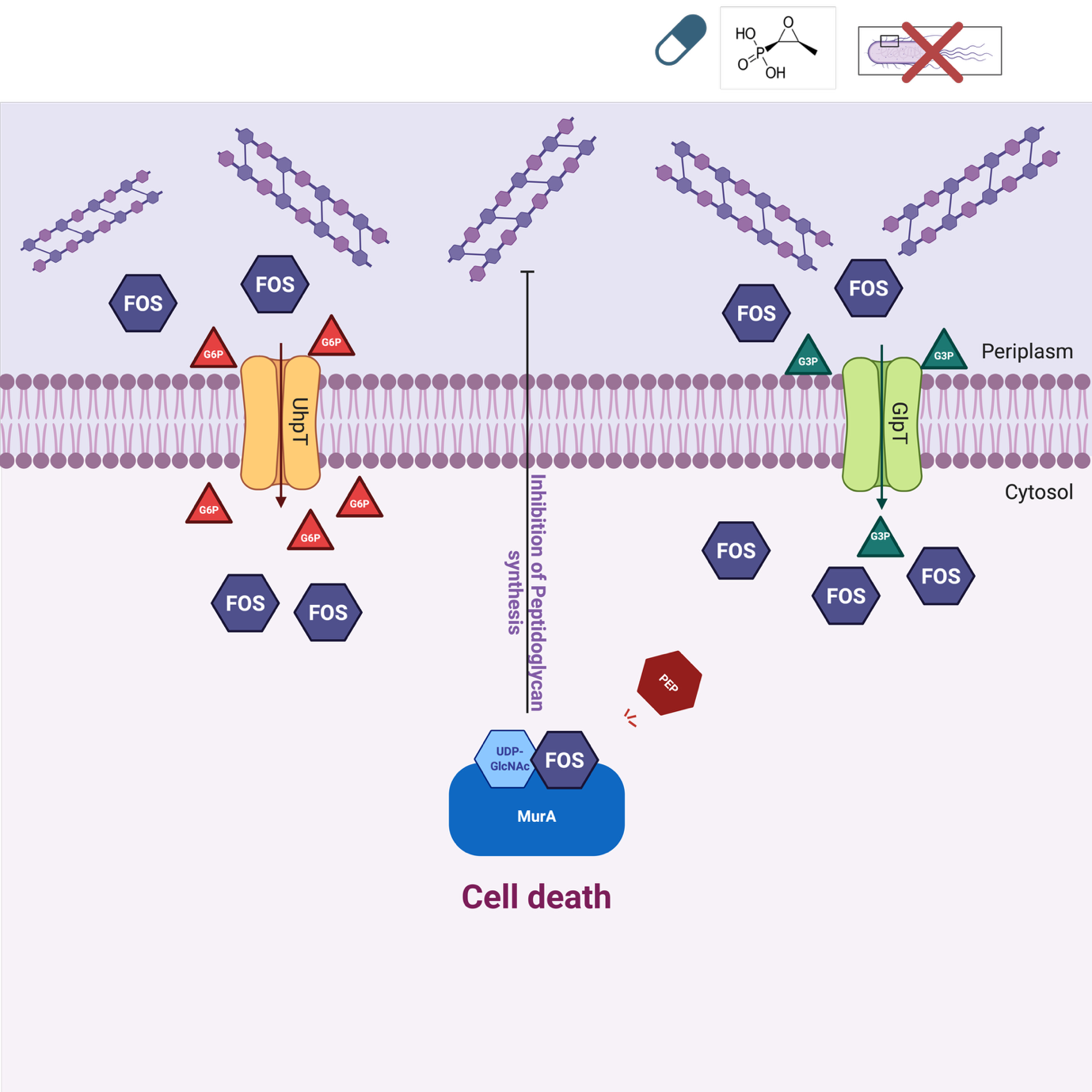

**Figure S2-:** FOS inhibition of peptidoglycan biosynthesis in bacteria.

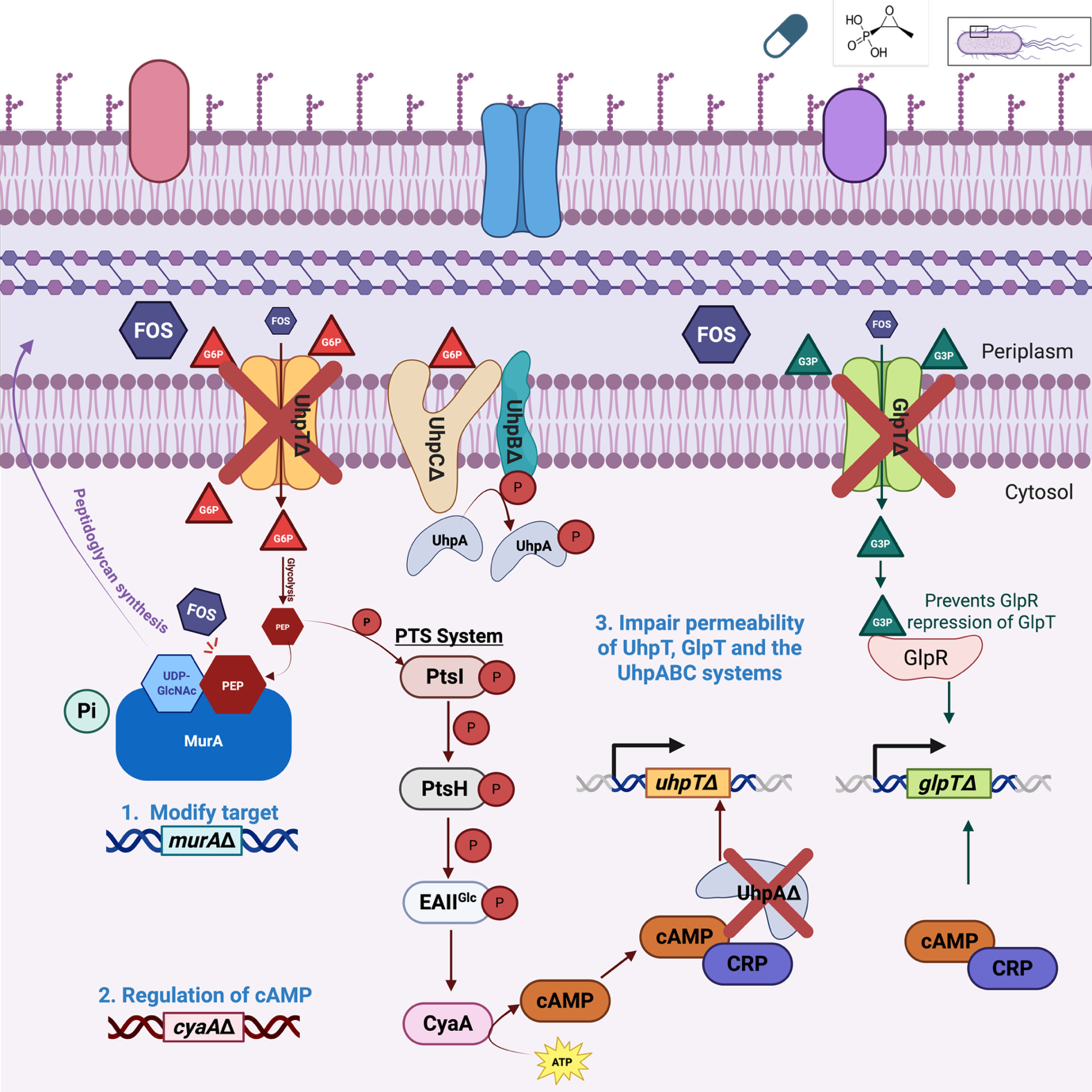

**Figure S3-:** Schematic illustration of known bacterial resistance mechanisms to FOS.

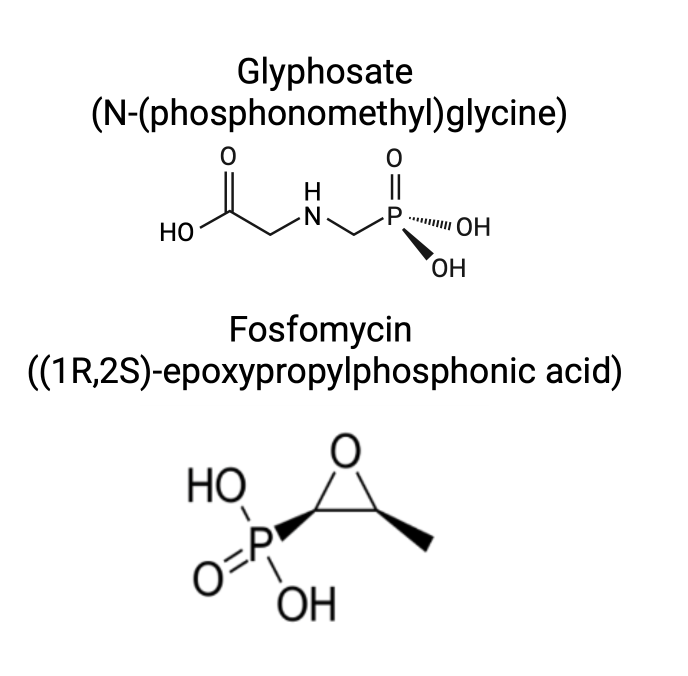

**Figure S4-:** Chemical structures of the phosphonates GLP and FOS.

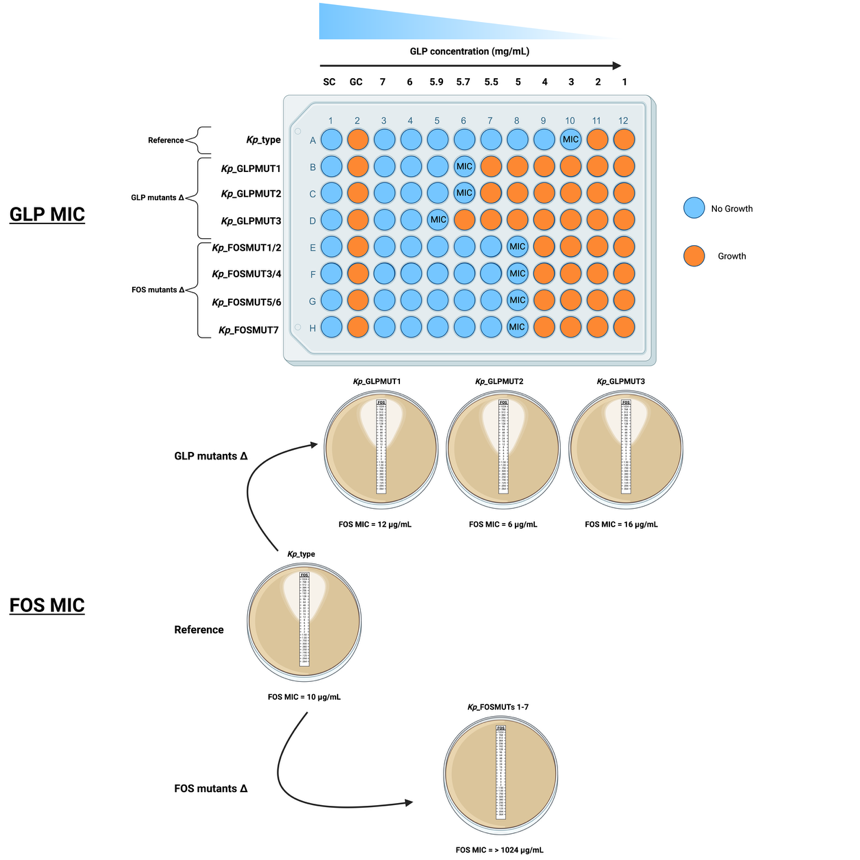

**Figure S5-:** Schematic of MIC determination for GLP and FOS. The workflow illustrates the methodology used to quantify MICs for the isogenic control (*Kp*_type) and its evolved derivatives, including three GLP-evolved (GLPMUT1-3) and seven FOS-evolved (FOSMUT1-7) mutants. GLP MICs were determined via the microbroth dilution method, while FOS MICs were determined using gradient diffusion strips (E-tests). SC = sterile control and GC = growth control.

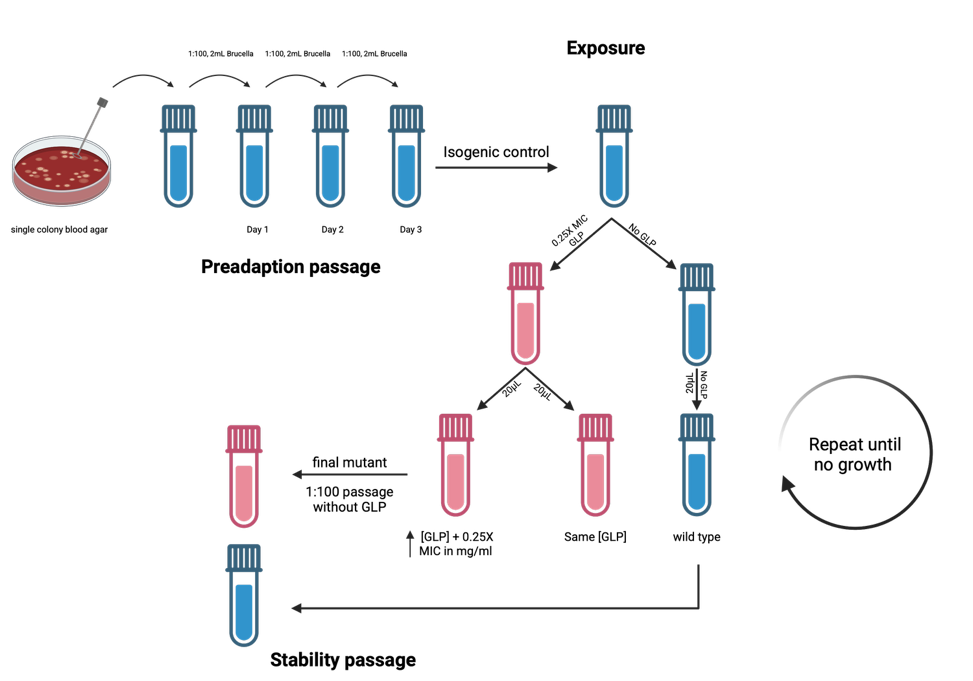

**Figure S6-:** Schematic figure of the steps involved in the selection of GLP and FOS mutants by adaptation to increasing concentrations of the herbicide.

**Table S1-:** Overview of the genome assembly data of the type strain and three GLP-evolved mutants.

| **Bacterial isolates**  **Features** | ***Kp*_type** | ***Kp*_GLPMUT1** | ***Kp*_GLPMUT2** | ***Kp*_GLPMUT3** |
| --- | --- | --- | --- | --- |
| Number of contigs | 120 | 86 | 145 | 93 |
| Number of contigs (>= 1,000 bp) | 88 | 64 | 121 | 64 |
| Total Length (bp) | 5571426 | 5434107 | 5429170 | 5430292 |
| Total Length (>= 1,000 bp) | 5549674 | 5419143 | 5412290 | 5410390 |
| Largest contig (bp) | 449065 | 500374 | 310373 | 501387 |
| GC (%) | 57.21 | 57.37 | 57.35 | 57.37 |
| N50^a^ | 225195 | 274590 | 95966 | 247487 |
| N90^b^ | 48597 | 82236 | 29628 | 77700 |
| L50^c^ | 9 | 8 | 20 | 8 |
| L90^d^ | 30 | 23 | 60 | 23 |
| Total CDS | 5180 | 5036 | 5145 | 5032 |
| rRNA | 14 | 14 | 12 | 14 |
| tRNA | 83 | 82 | 83 | 82 |

^a^ N50 is defined as the length of the contig which, along with all the longer contigs, covers at least 50% of the whole genome.

^b^ N90 is defined as the length of the contig which, along with all the longer contigs, covers at least 90% of the whole genome.

^c^ L50 is defined as the lowest number of contigs required to cover at least 50% of the whole genome.

^d^ L90 is defined as the lowest number of contigs required to cover at least 90% of the whole genome.

**Table S2-:** Overview of the genome assembly data of the seven FOS- evolved mutants.

| **Bacterial isolates**  **Features** | ***Kp*_FOSMUT1** | ***Kp*_ FOSMUT2** | ***Kp*_ FOSMUT3** | ***Kp*_ FOSMUT4** | ***Kp*_FOSMUT5** | ***Kp*_ FOSMUT6** | ***Kp*_ FOSMUT7** |
| --- | --- | --- | --- | --- | --- | --- | --- |
| Number of contigs | 135 | 127 | 287 | 130 | 117 | 114 | 128 |
| Number of contigs (>= 1,000 bp) | 101 | 99 | 243 | 100 | 91 | 92 | 103 |
| Total Length (bp) | 5547349 | 5547942 | 5536135 | 5540717 | 5549639 | 5539837 | 5541780 |
| Total Length (>= 1,000 bp) | 5524647 | 5528726 | 5506000 | 5520082 | 5531912 | 5524685 | 5525133 |
| Largest contig (bp) | 448759 | 449142 | 138128 | 382529 | 449325 | 448501 | 382401 |
| GC (%) | 57.21 | 57.22 | 57.2 | 57.22 | 57.22 | 57.23 | 57.22 |
| N50^a^ | 169930 | 179627 | 43121 | 152520 | 225996 | 178509 | 152520 |
| N90^b^ | 41972 | 43827 | 11253 | 41953 | 50772 | 43445 | 42182 |
| L50^c^ | 11 | 11 | 42 | 12 | 9 | 10 | 12 |
| L90^d^ | 35 | 33 | 137 | 37 | 30 | 34 | 37 |
| Total CDS | 5160 | 5160 | 5342 | 5153 | 5164 | 5155 | 5152 |
| rRNA | 9 | 10 | 6 | 5 | 10 | 5 | 5 |
| tRNA | 82 | 82 | 79 | 85 | 81 | 84 | 84 |

**Table S3-:** The number of SNPs identified after WGS analysis and corresponding FOS MIC values for a collection of FOS-evolved Klebsiella mutants.

| ***Klebsiella*** | **No. of SNPs** | **FOS (μg/ml)** |
| --- | --- | --- |
| ***Kp*_type** | **-** | **80** |
| *Kp*_FOSMUT1 | 11 | 160 |
| *Kp*_FOSMUT2 | 14 | 200 |
| ***Kp*_FOSMUT3** | **7,317** | **240** |
| *Kp*_FOSMUT4 | 22 | 280 |
| *Kp*_FOSMUT5 | 13 | 320 |
| *Kp*_FOSMUT6 | 14 | 360 |
| *Kp*_FOSMUT7 | 21 | 400 |

**Table S4-:** Summary of the pangenome of the isogenic *Kp*_type strain and its three GLP-evolved and seven FOS-evolved mutants.

| **Pangenome** | **% strains** | **Number of genes** |
| --- | --- | --- |
| Core genes | (99% <= strains <= 100%) | 4897 |
| Soft core genes | (95% <= strains < 99%) | 0 |
| Shell genes | (15% <= strains < 95%) | 366 |
| Cloud genes | (0% <= strains < 15%) | 364 |
| Total genes | (0% <= strains <= 100%) | 5,627 |

**Table S5-:** Gene name, locus tag, and product annotation of hypermutator phenotypes as obtained from NCBI (Reference Sequence NC_009648.1). GM refers to GLP mutant and FM refers to FOS mutant.

| **Gene name** | **Locus tag** | **Product annotation** | | **SNP position** |
| --- | --- | --- | --- | --- |
| ***mutS*** | KPN_03094 | DNA mismatch repair protein | GM2= Gln476* ; Pro235Pro ; 632_633delAGinsTA p.Gln211Leu  FM3= Ile279Val ; Ser89Ser | |
| ***mutL*** | KPN_04568 | DNA mismatch repair protein | GM2= Gly515Glu ; Gly479Gly ; Lys307Thr  FM3= His308Tyr | |
| ***mutH*** | KPN_03240 | DNA mismatch repair protein | GM2= Leu78Leu | |
| ***uvrD*** | KPN_04312 | DNA helicase II | GM2= Asp250Glu ; Asp311Ala ; Glu447*  FM3= Val18Val ; Arg3775His ; Arg619Arg ; Val720Gly | |
| ***dnaQ*** | KPN_00230 | DNA polymerase III subunit epsilon | GM2= Lys32* ; Ser160Ser ; Ser241Thr | |
| ***mutY*** | KPN_03393 | Adenine DNA glycosylase | GM2= His102Gln ; Trp26Arg | |
| ***mutT*** | KPN_00103 | 8-oxo-dGTP diphosphatase | GM2= Val62Val  FM3= Val8Asp ; Pro36leu | |

**Table S6-:** Gene name, locus tag, and product annotation of shared SNPs as obtained from NCBI (Reference Sequence NC_009648.1) and *EnsemblBacteria* for *Klebsiella pneumoniae subsp. pneumoniae* strain MGH 78578 (GCA_000016305). GM refers to GLP mutant and FM refers to FOS mutant.

| **Gene name** | **Locus tag** | **Product annotation** | | **SNP position** |
| --- | --- | --- | --- | --- |
| ***rnfC*** | KPN_01966 | electron transport complex protein RnfC/Ion-translocating oxidoreductase complex subunit | FM6+FM7= Ala561Ala  FM7=Asp578Glu;Ala599Ala;Ala604Ala  FM3=Glu559Ala;Ala561Ala;Glu627Ala  GM2=Thr293Ala;Thr345Thr;Gln543Leu | |
| ***traC*** | KPN_pKPN4p07139 | F pilus assembly protein | FM4=Ser758Ser | |
| **KPN_04348/23S** | KPN_04348 | 23S ribosomal RNA (partial) | FM4=rRNA intergenic region | |
| ***nrdE*** | KPN_03006 | class 1b ribonucleoside-diphosphate reductase subunit alpha | GM3=Ala253Ala  GM2=Asp293Glu;Glu123Glu;Lys35Arg  FM3=Val500Ala;Ala299Ala;His259Arg;  Gln142Gln;Glu68Gly;Arg37Gly | |
| ***nemA*** | KPN_01989 | N-ethylmaleimide reductase/ alkene reductase | GM3=Asp253Ala  GM2=Arg230Arg  FM3=Ser147* | |
| ***rhmD*** | KPN_02652 | L-rhamnonate dehydratase | GM3=Ala207Glu  GM2=Pro398Leu  FM3=Ile199Ile;Tyr28Ser | |
| ***yjeH*** | KPN_04531 | L-methionine/branched-chain amino acid transporter | GM3=Asp119Asp  GM2=Phe110Cys;Pro170Leu;  Glu209Lys;Leu227Phe  FM3=Arg215Arg;Leu336Phe | |
| ***bioC*** | KPN_00802 | malonyl-ACP O-methyltransferase BioC/ biotin biosynthesis; reaction prior to pimeloyl CoA | GM3=Ala96Ala  GM2=Arg36Gly  FM3=Glu148Gly;Ala63Glu | |
| ***topB*** | KPN_01208 | DNA topoisomerase III | GM3=Val182Val  GM2=Ala606Ser;His549Asn;  His340Tyr;Glu7Gly | |
| ***dcuC*** | KPN_00654 | anaerobic C4-dicarboxylate transporter DcuC/ dicarboxylate transport protein (*DcuC* family) | GM3=Glu427Asp  FM3=Leu274His;Leu360Met | |
| ***leuD*** | KPN_00077 | 3-isopropylmalate dehydratase small subunit/  isopropylmalate isomerase small subunit | GM3=Ile164leu  GM2=Ile189Thr;Asn174Thr;Leu148Leu  FM3=Pro66Arg | |
| **f*imD*** | KPN_01671 | putative fimbrial biogenesis outer membrane usher protein | GM3=Gln347Pro  GM2=Ser541Cys;Asp473Glu;  Tyr397Cys;Tyr345Tyr;Tyr172His  FM3=Leu825Val;Pro800Pro;  Val747Val;Ser583Cys | |
| ***hpxK*** | KPN_01761 | allantoate amidohydrolase/peptidase | GM3=Ser358Cys;Ile370Ser  GM2=Gly210Ala;Val256Ala;Tyr332*  FM3=Thr275Thr;Ala345Pro | |
| ***oxlT*** | KPN_01750 | Oxalate/formate MFS antiporter | GM3=Ile193Thr  GM2=Ala146Ala;Ser258Leu  FM3=Val244Phe | |
| **KPN_04884** | KPN_04884 | putative prophage protein | GM3=Phe34Phe | |
| ***czcO*** | KPN_01462 | flavin-containing monooxygenase/ FAD-dependent oxidoreductase | GM3=His156Pro  FM3=Gly380Gly;Trp352Cys;Pro65Ala;Ala42Val | |
| **KPN_pKPN3p05980** | KPN_pKPN3p05980 | Hypothetical protein | FM6+FM7= Arg16Ser | |
| **KPN_pKPN5p08171** | KPN_pKPN5p08171 | putative antirestriction protein | GM3+FM4+FM7=Ser15Cys;  GM3+FM1+FM2+FM4+FM7=Glu13Glu | |
| **KPN_pKPN5p08175** | KPN_pKPN5p08175 | DNA methylase/ Methyltransferase | FM5= Asp83Ala | |
| **KPN_pKPN5p08222** | KPN_pKPN5p08222 | Hypothetical protein | GM1+GM2+GM3+FM1+FM2+FM3  +FM4+FM5+FM6+FM7 = Gly115Gly  GM2=Gln169Leu;Pro112Arg;Asp110Glu;Phe108Ser.  Gly67Gly;Thr61Met  FM3=Val135Ala;Phe96Leu | |
